# A computational framework for a Lyapunov-enabled analysis of biochemical reaction networks

**DOI:** 10.1101/696716

**Authors:** M. Ali Al-Radhawi, David Angeli, Eduardo D. Sontag

## Abstract

Complex molecular biological processes such as transcription and translation, signal transduction, post-translational modification cascades, and metabolic pathways can be described in principle by biochemical reactions that explicitly take into account the sophisticated network of chemical interactions regulating cell life. The ability to deduce the possible qualitative behaviors of such networks from a set of reactions is a central objective and an ongoing challenge in the field of systems biology. Unfortunately, the construction of complete mathematical models is often hindered by a pervasive problem: despite the wealth of qualitative graphical knowledge about network interactions, the form of the governing nonlinearities and/or the values of kinetic constants are hard to uncover experimentally. The kinetics can also change with environmental variations.

This work addresses the following question: given a set of reactions and without assuming a particular form for the kinetics, what can we say about the asymptotic behavior of the network? Specifically, it introduces a class of networks that are “structurally (mono) attractive” meaning that they are incapable of exhibiting multiple steady states, oscillation, or chaos by virtue of their reaction graphs. These networks are characterized by the existence of a universal energy-like function called a *Robust Lyapunov function* (RLF). To find such functions, a finite set of rank-one linear systems is introduced, which form the extremals of a linear convex cone. The problem is then reduced to that of finding a common Lyapunov function for this set of extremals. Based on this characterization, a computational package, Lyapunov-Enabled Analysis of Reaction Networks (`LEARN`), is provided that constructs such functions or rules out their existence.

An extensive study of biochemical networks demonstrates that `LEARN` offers a new unified framework. Basic motifs, three-body binding, and genetic networks are studied first. The work then focuses on cellular signalling networks including various post-translational modification cascades, phosphotransfer and phosphorelay networks, T-cell kinetic proofreading, and ERK signalling. The Ribosome Flow Model is also studied.

**Author summary:** A theoretical and computational framework is developed for the identification of biochemical networks that are “structurally attractive”. This means that they only allow global point attractors and they cannot exhibit any other asymptotic behavior such as multi-stability, oscillations, or chaos for any choice of the kinetics. They are characterized by the existence of energy-like functions. A computational package is made available for usage by a wider community. Many relevant networks in molecular biology satisfy the assumptions, and some are analyzed for the first time.

## Introduction

Many biological systems are known for the ability to operate precisely and consistently subject to potentially large disruptions and uncertainties [1–5]. Examples are homeostasis, understood as the maintenance of a desired steady state (perhaps associated to an observable phenotype) against the variability of in-vivo concentrations of biochemical species, or a consistent dynamical behavior in the face of environmental variations which change the speed of reactions.

The vaguely defined term “robustness” is often used to refer to this consistency of behavior under perturbations. The present work deals with such notions of “biological robustness”, as well with a “robustness of analysis” notion in which conclusions can be drawn despite inaccurate mathematical models.

Models of core processes in cells are typically biochemical reaction networks. This includes binding and unbinding, production and decay of proteins, regulation of transcription and translation, metabolic pathways, and signal transduction [6]. However, in contrast to engineered chemical systems, biology poses particular challenges. On the one hand, the reactants and the products in such interactions are frequently known, and hence the *species-reaction graph* is available. On the other hand, the exact form and parameters (i.e, kinetics) that determine the speed of transformation of reactants into products are often unknown. This lack of information is a barrier to the construction of complete mathematical models of biochemical dynamics. Even if the kinetics are exactly known at a specific point in time, they are influenced by environmental factors and hence they can change. Hence, the ability to draw conclusions regarding the qualitative behavior of such networks without knowledge of their kinetics is highly relevant, and has been advocated under the banner of “complex biology without parameters” [4]. But is such a goal realistic? It is known that the long-term qualitative behavior of a nonlinear system can be critically dependent on parameters, a phenomenon known as bifurcation. This fundamental difficulty led to statements such as Glass and Kauffman’s 1973 assertion that “it has proved impossible to develop general techniques which may be applied to find the asymptotic behavior of complex chemical systems” [7].

Notwithstanding such difficulties, many classes of reaction networks are observed to have a “well-behaved” qualitative long-term dynamical behavior for wide ranges of parameters and various types of nonlinearities. This means specifically in our context that such networks do not have the potential for exhibiting complex steady-state phenotypes such as multiple-steady states (e.g, toggle switches), oscillations (e.g, repressilator), or chaos. Their typical behavior is that the concentrations eventually settle into a unique steady steady (called an *attractor*) for any initial condition (with fixed *total* substrate, gene and enzyme concentrations). Hence, we call them *structurally attractive*. The relevant biological phenotype for such networks is the unique attractor, which is mathematically represented by the concentrations of the biochemical species at steady state. Discerning such networks is not generally trivial. For instance, within the class of post-translational modification (PTM) cycles, some cascades are “structurally attractive” but others can exhibit oscillations and multistability [8]. Figure 1 illustrates the typical behavior of an attractive network vs a multistable network for two PTM cycles that have been proposed as models for double phosphorylation. We will study PTM cycles in detail later in the paper.

**Fig 1.**
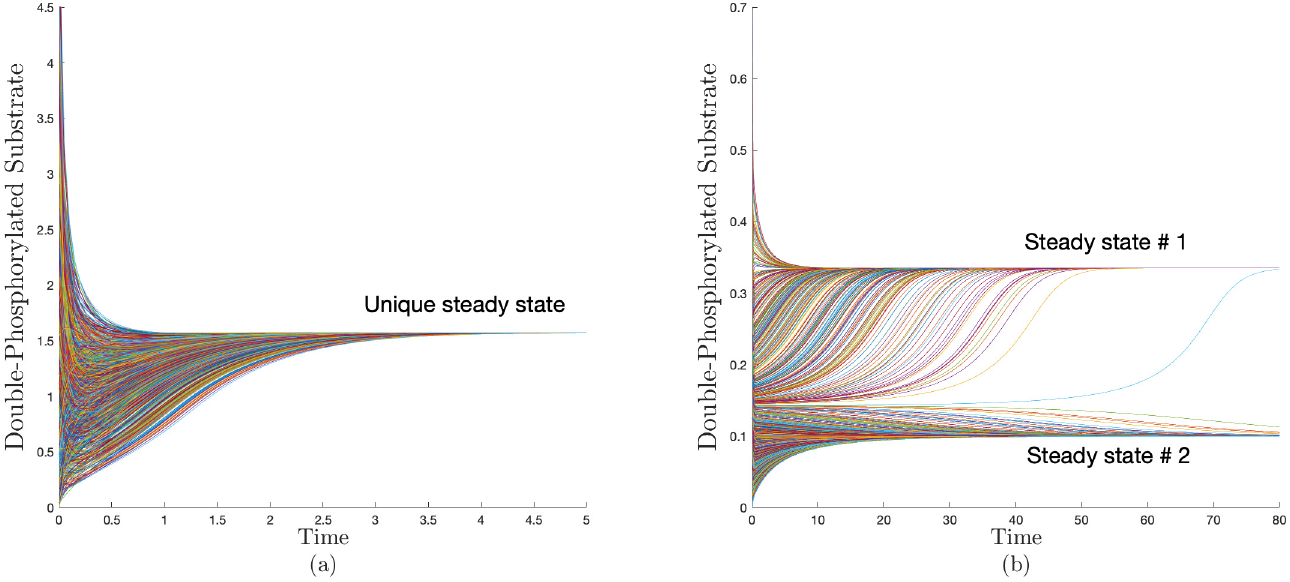
Distinct qualitative behaviors for two models of a double PTM. This is illustrated by the time series plots for the double phosphorylated substrate with randomized initial conditions for fixed total substrate and enzyme concentrations. (a) the processive mechanism exhibits a unique global attractor, (b) a distributive mechanism exhibits multistability for some parameters. See Figure 8 and the accompanying discussion. The parameters are given in S1 Text-§6.

In the terminology of dynamical and control system theories, the defining feature of an attractive network is that it can only exhibit global point attractors (i.e., unique globally asymptotically stable steady states). The classical way to certify stability is by exhibiting an appropriate energy-like function, commonly referred to as a *Lyapunov function* [9, 10]. Existence of such a function provides many guarantees on qualitative behavior, including notably the fact that its sub-level sets act as trapping sets for trajectories [11]. Furthermore, they allow the development of a systematic study of model uncertainties and response to disturbances [9, 10]. However, it is notoriously difficult to find such functions for nonlinear systems due to the lack of general constructive techniques.

The search of Lyapunov functions for nonlinear reaction networks can be traced back to Boltzmann’s *H*-Theorem [12], which applies only to the restrictive subclass of detailed-balanced networks. Wei [13] in 1962 postulated that all chemical systems should satisfy an “axiom of convergence” and there shall exist a suitable Lyapunov function. Perhaps the most striking success in this line of thought was the development of the Horn-Jackson-Feinberg (HJF) theory of complex-balanced networks [14–17] in the early 1970s, which relies on using the sum of all the chemical *pseudo*-energies stored in species as a Lyapunov function. When specific graphical conditions are satisfied, complex-balancing is guaranteed for all kinetic constants. Global stability can be proven in certain cases [18, 19]. Despite the elegance and theoretical appeal of the method, the assumptions needed for its applicability are restrictive, and are not widely satisfied in biological models. For example, many basic motifs (e.g, transcription/translation and enzymatic reactions) are not complex balanced. Furthermore, HJF theory assumes, although with some exceptions, that the reaction kinetics are Mass-Action. It has been argued that this assumption “is not based on fundamental laws” and is merely “good phenomenology” [20]. These laws are usually justified by the intuitive image of colliding molecules. However, this is often not the right level of analysis for biological modeling, where alternative kinetics such as Michaelis-Menten and Hill kinetics are used in situations involving multiple time scales [21].

Beside complex-balanced networks, a few additional classes of attractive networks have been identified. These include mono-molecular networks, which can be handled within the framework of compartmental systems using a Lyapunov function [22, 23]. More recently, global convergence has been shown for another class of networks via the concept of monotonicity without supplying a Lyapunov function [24] where sufficient graphical conditions have been given.

In previous work [25–27], two of the authors proposed a direct approach to the problem, introducing the class of piecewise linear-in-rates functions, which act as Lyapunov functions regardless of the specific form of the reaction nonlinearities or kinetic constants. They guarantee the uniqueness of steady states and global stability under mild additional conditions.

In this work, the results from [25–27] are generalized in several directions, theoretically, computationally, and in terms of biological applications. First, we propose a general characterization of “structurally attractive” networks. We require the existence of a universal rate-dependent function, which we call a Robust Lyapunov Function (RLF), that is a Lyapunov function for any choice of the kinetics. We proceed to propose a computational framework for finding such functions. To this end, the dynamics of the network are embedded in a linear convex cone. The *extremals* of this cone are a set of rank-one matrices that derive from the stoichiometry of the network. If a common Lyapunov function exists for the extremals, then it can be used to construct an RLF and the network is certified to be *attractive*. In the special case that kinetics are mono- or bimolecular, the RLF is piecewise linear or piecewise quadratic on species, respectively.

Computationally, we complement previous reaction network toolboxes [28, 29] and we provide a Lyapunov-Enabled Analysis of Reaction Networks (`LEARN`) toolbox to implement the results on any given network by searching for an RLF and checking the appropriate conditions via four main methods: a graphical algorithm, a linear program, an iterative procedure, and a semi-definite program. Additionally, `LEARN` checks for conditions that rule out the existence of an RLF.

We then proceed to carry out an extensive discussion of biochemical networks to show the applicability of our results. Foundational studies in systems biology [6] have revealed that biochemical networks have many common “motifs”. We show that our results form a basis for the understanding of the behavior of a large class of networks of various degrees of complexity. They may be applied to study basic motifs such as binding/unbinding, three-body binding, transcription and translation networks, and enzymatic reactions. Most cellular signalling involves PTMs as building blocks, and their malfunction is frequent in diseases such as cancer and Alzheimer [30, 31]. Hence, we study in details PTMs cascades, ERK signalling, and phosphotransfer and phosphorelay networks. In addition, we study important biological networks such as T-cell kinetic proofreading, and the Ribosome Flow Model. We show that our Lyapunov functions can be used to construct safety sets and perform dynamic flux analysis. Many of the networks studied are not amenable to the previously-mentioned analysis techniques, HJF theory in particular. A comparison with other methods in included in the Discussion (see Table 1). In particular, our results include the class of monomolecular networks treated in [22, 23], and it applies all the biochemical networks studied in [32], [24], [33]. A preliminary version of a subset of these results were presented in conferences [34], [35].

**Table 1.**
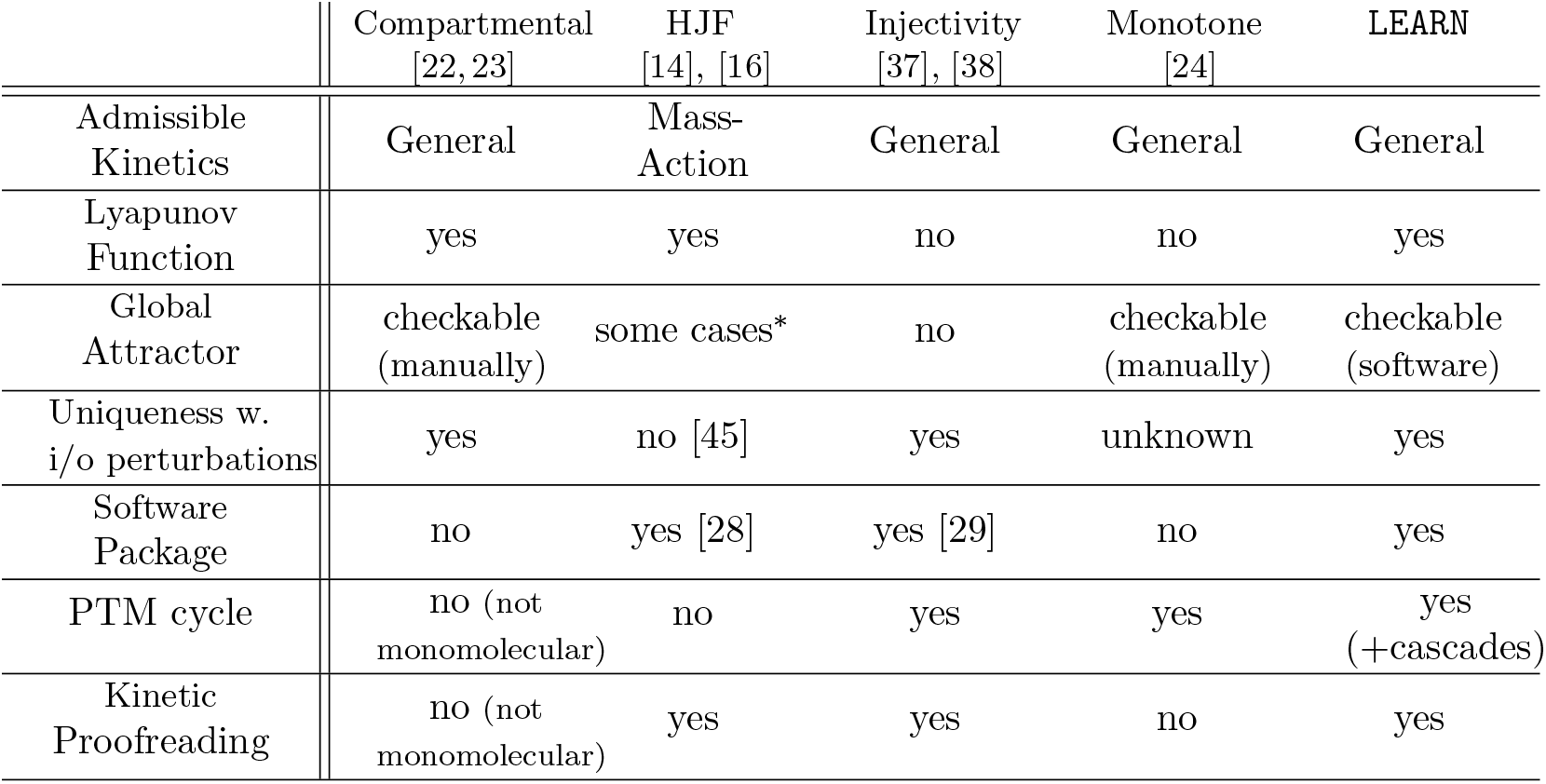
Comparison with other methods in the literature. The row that corresponds to “admissible kinetics” asks about the functional form of the reaction rates for which the method is applicable. “Global attractor” asks whether the method is able to provide guarantees for the global convergence to an attractor. “Uniqueness with i/o perturbations” asks whether the method can guarantee uniqueness of steady states with respect to arbitrary addition of inflows and outflows to the network (i.e., “homogeneous CFSTR” in the terminology of [45]). Rows that correspond to “PTM cycle” and “Kinetic proofreading” ask whether the method can tackle the networks in Figure 2 and Figure 10, respectively. We have picked these two networks as non-trivial examples that are pertinent to systems biology. The question of a global attractor for HJF-type networks is marked by an asterisk (*) since a proof has been proposed in a preprint [46] but is not formally published yet. (See [47] also)

|  | Compartmental<br>[22, 23] | HJF<br>[14], [16] | Injectivity<br>[37], [38] | Monotone<br>[24] | LEARN |
| --- | --- | --- | --- | --- | --- |
| Admissible Kinetics | General | Mass-Action | General | General | General |
| Lyapunov Function | yes | yes | no | no | yes |
| Global Attractor | checkable (manually) | some cases* | no | checkable (manually) | checkable (software) |
| Uniqueness w. i/o perturbations | yes | no [45] | yes | unknown | yes |
| Software Package | no | yes [28] | yes [29] | no | yes |
| PTM cycle | no (not monomolecular) | no | yes | yes | yes (+cascades) |
| Kinetic Proofreading | no (not monomolecular) | yes | yes | no | yes |

Theoretically, our results connect with a corpus of previous literature. We show that the RLFs can be formulated in different coordinates, and how this relates to the ones proposed in [34], [36]. Also, the approach makes contact with the notions of structural injectivity [37–40], structural persistence [41], and uncertain linear systems [42–44].

### Overview and Comparison

The paper has been written for a diverse readership, and has been structured accordingly. Readers who are interested in the general concepts, the biological applications, and the software package only need to consult the Introduction, the Results, and `LEARN`’s accompanying manual (SI §7). Users can apply the results by supplying the list of reactions encoded as a stoichiometry matrix as an input to `LEARN`’s main subroutine for a report of results. Readers who are also interested in the technical mathematical details can consult the Methods section.

Since `LEARN` guarantees that a certain mechanism cannot admit multistability, oscillation, or chaos, it can be used to distinguish competing biochemical reaction networks at the modeling step. We give an example of this when discussing processive vs distributive post-translational cycles.

`LEARN` can be compared to other results in the literature as shown in Table 1.

### Terminology and Motivational Example

A list of reactions can be abstracted mathematically into the framework of Chemical Reaction Networks (CRNs). A CRN consists of a set of species 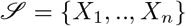 and a set of reactions 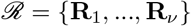. (see Methods for an elaborate discussion) Figure 2-a) gives an example of a reaction network for a core signaling motif which is the standard post-translational modification (PTM) cycle [48, 49]. The relative gain or loss of molecules of each species in a reaction is encoded in a matrix Γ ∈ ℝ^*n*×*ν*^ called *the stoichiometry matrix*. It is given in Figure 2-b for the PTM cycle. CRNs admit graphical representations naturally. A CRN can be modeled as a graph with two sets of nodes: reactions and species. Mathematically, it is a bipartite weighted directed graph, called *the species-reaction graph* (or a Petri-net [50]). The graph corresponding to the PTM cycle is given in Figure 2-c). The stoichiometry matrix Γ becomes the *incidence matrix* of the graph [51].

**Fig 2.**
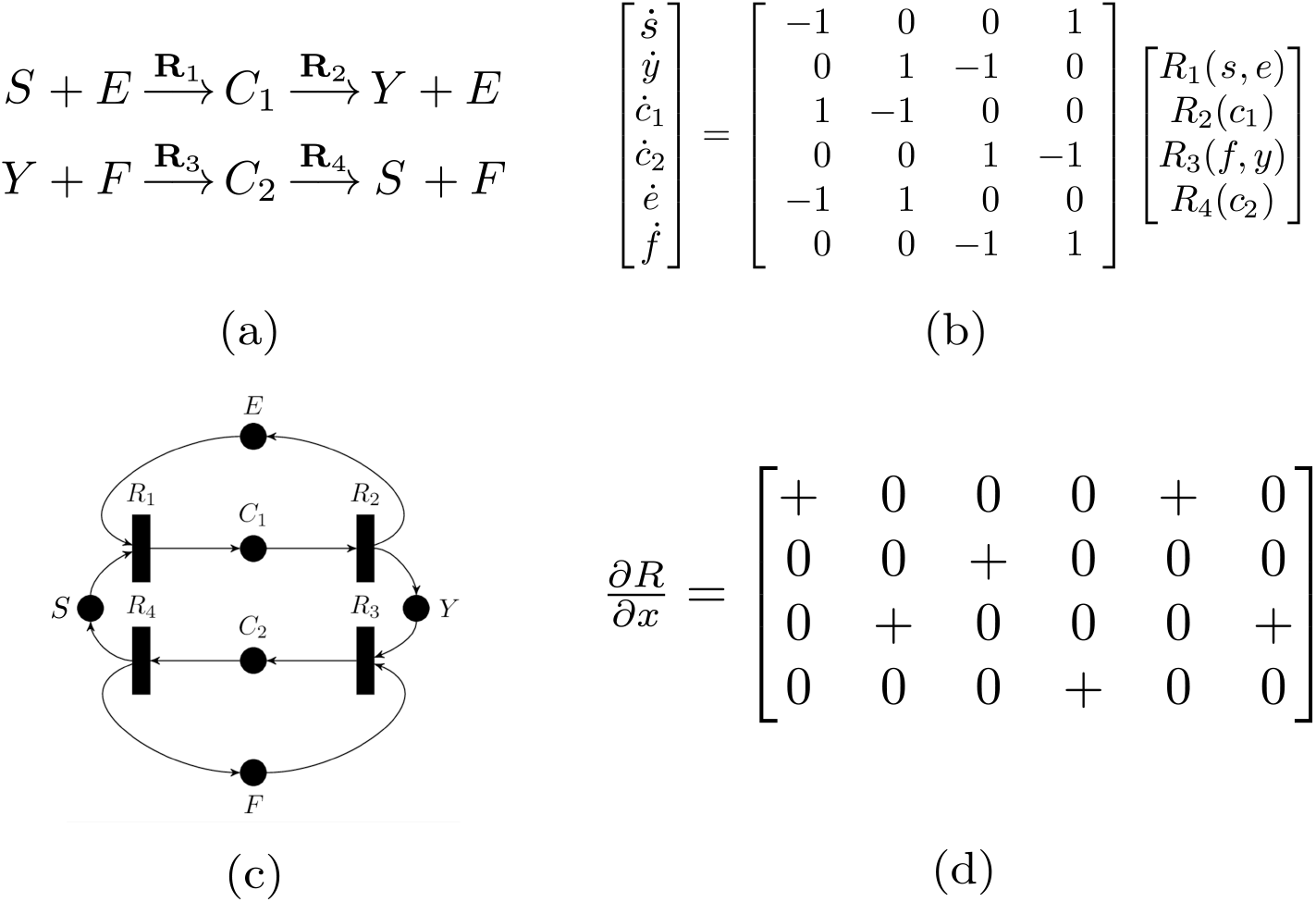
Illustration of a post-translational modification reaction network. (a) The list of reactions with six species. A kinase *E* interacts with a substrate *S* to form a complex *C*_1_ which transforms into a phosphorylated substrate *Y*. Similarly, a phosphatase *F* dephosphorylates *Y* back to *S* via an intermediate complex *C*_2_. (b) The ODE equation description of the time-evolution of the concentration of the species. (c) The graphical representation of the network as a Petri-net. A circle represents a species and a rectangle represents a reaction, (d) The Jacobian matrix of the reaction rate vector. This is the only information we assume to be known about *R*(*x*).

As we are interested in studying the long-term dynamical behavior, a concentration *x_i_* ≥ 0, *i* = 1, ‥, *n* is assigned to each species. Hence, the concentration vector at time *t* is *x*(*t*) = [*x*_1_(*t*), …, *x_n_*(*t*)]^*T*^. A reaction rate (or flux) *R_j_*(*x*), *j* = 1, ‥, *ν* is assigned to each reaction. The reaction rate vector is *R*(*x*) = [*R*_1_(*x*), …, *R_ν_*(*x*)]^*T*^. The time-evolution of the concentration vector is given by the standard ordinary differential equation (ODE) given as [52]:

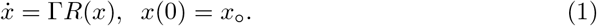

Biochemical networks usually contain conserved quantities (i.e., moieties) such as the total amount of enzymes, substrates, ribosomes, RNA polymerase, etc. For each conserved quantity, there there exists a nonnegative vector *d* such that *d^T^*Γ = 0, and *d* is called a conservation law. If every species is supported in at least one conservation law the network is said to be *conservative*. For example, the PTM cycle in Figure 2 is conservative with three conservation laws *c*_1_ + *c*_2_ + *x* + *y* = [*X*]_*total*_, *e* + *c*_1_ = [*E*]_*total*_, and *f* + *c*_2_ = [*F*]_*total*_, which are the total amounts of the substrate and the two enzymes, respectively, and they stay constant throughout the reaction. Hence, claims of global stability and uniqueness of steady states are relative to the conserved quantities. A set of concentrations that shares the same conserved quantities is called a *stoichiometric class*.

For the PTM cycle, the ODE is given in Figure 2-b). We do *not* assume that the reaction rates have a specific functional form such as Mass-Action. We only assume that the rates are *monotone*, meaning that as the concentration of reactants increases, the rate of the reaction increases (see Methods). This can be interpreted as enforcing a specific sign pattern on the partial derivatives of *R*. This means that all the entries of the *Jacobian* matrix of *R* (i.e, *∂R/∂x*), are either zero or non-negative. For the PTM cycle, Figure 2-d) illustrates our assumptions on the reaction rates encoded in terms of the Jacobian matrix. Such reactions include all common reaction rates such as Mass-Action, Michaelis-Menten, Hill, etc.

Despite its application relevance, establishing the long-term behavior of the PTM cycle in Figure 2 was an open problem till the 2000s. HJF’s theory cannot be used for deciding stability since the PTM cycle is a non-zero deficiency network. In 2008, this problem was tackled via monotonicity techniques [24, 32], but no Lyapunov function has been provided. As a motivation, we study the same cycle using our proposed method. An intuitive way to approach its analysis is to consider the central loop in Figure 2, and then study the sum of absolute rate differences along it. This can be loosely motivated by considering the reactions rates as potentials and the concentration of species as charges, and noting that the difference of “potentials” causes the concentration of species to change via the flow of a “current”. Hence, we define the *i*th current as the rate of change of the concentration of the *i*th species. Thus, we consider the weighted *sum of currents* 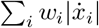 as a candidate Lyapunov function. It can also be written as follows:

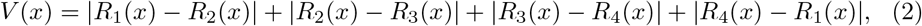

which is a piecewise linear-in-rates function. In order to verify whether this is indeed a Lyapunov function, we can analyze it region-wise to check that it decreases along trajectories. Consider for instance the region 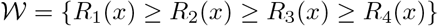. The candidate *V* simplifies to the difference of “potentials” across the substrate *S*:

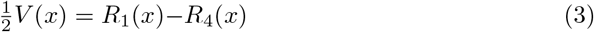

To evaluate 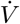, we need the signs of the “currents” 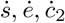. In our example, we can use the inequalities defining 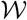 so that the signs can be read from the graph as follows: 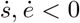 and 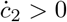. By noting that these signs are matched to the coefficients of *R*(*x*) in (3), and since *∂R/∂x* is nonnegative, we can write the following inequality in 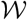:

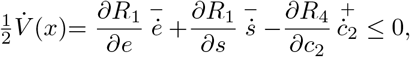

where the sign of the rate of change of each concentration is indicated above it.

Therefore, 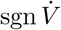 can be determined conclusively *without* knowing the kinetics. In fact, this can be repeated for all regions to conclude that *V* is non-increasing along all possible trajectories of (1). (See the Results section for further analysis)

The lesson that can be drawn from this example is that a robust analysis of reaction networks can be carried out by considering candidate Lyapunov functions of the form 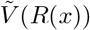 that vanish exactly on the steady state set, i.e. the set {*x*|Γ*R*(*x*) = 0}. This approach does not require the computation of the actual steady state.

### Robust Lyapunov Functions

The motivating example has shown that we can have a Lyapunov function 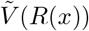 that decreases along trajectories for any monotone kinetics *R*. Hence for a given network 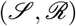 we will be looking for a function 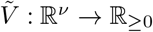 that vanishes only on the set of steady states, i.e

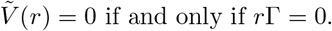

Furthermore, 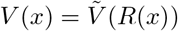 needs to be nonincreasing along the trajectories of (1), i.e it must satisfy:

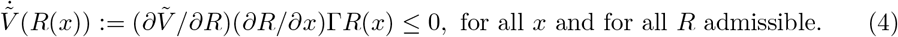

If such a function exists then we call it a *Robust Lyapunov Function* (RLF), and the network is called *structurally attractive*. Mathematically, the RLF needs only to be locally Lipschitz and the derivative is defined in the sense of Dini’s (see Methods).

#### Example (cont’d)

For the PTM cycle (Figure 2) the function 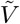 is 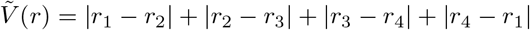.

## Results

### Characterization of RLFs

The above definition of an RLF does not offer a constructive way for finding one or for checking a candidate. Our first result is to give a characterization of RLF in terms of a set of rank-one linear systems, each of which corresponds to a *reaction-reactant pair*. The set of all such pairs is 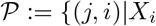 participates in the reaction R_j_}. Let *s* be total number of such pairs. Then, 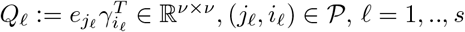 where {*γ*_1_, ‥, *γ_n_*} are the rows of Γ and {*e*_1_, ‥, *e_ν_*} are columns of the *ν* × *ν* identity matrix.

The matrices *Q*_1_, ‥, *Q_s_* will serve as system matrices for *s* linear systems and also as extremals of a linear convex cone. We show (see Methods) that (*∂R/∂x*)Γ ∈ cone(*Q*_1_, ‥, *Q_s_*) = {∑_*ℓ*_*ρ*_*ℓ*_*Q*_*ℓ*_|*ρ*_*ℓ*_ ≥ 0}. We will be looking for a function 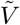 that acts as a *common Lyapunov function* for these linear systems and satisfies 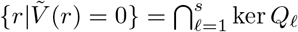 (see Methods).

#### Example (cont’d)

For the PTM cycle (Figure 2), the extremals are 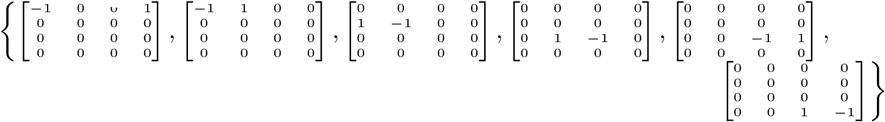.

We are ready to state the main result of this section. (See Methods)

#### Theorem 1.

*Given* 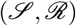. *Let* (1) *be the associated ODE. A function* 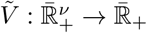 *is a common Lyapunov function for the set of linear systems* 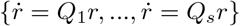 *if and only if* 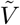 *is an RLF for the reaction network* 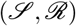.

### The Search for RLFs

The characterization provided in Theorem 1 can be used for devising computational algorithms that search for an RLF. In Methods, we present several algorithms for constructing piecewise linear (PWL) or piecewise quadratic RLFs. In order to simplify the presentation, we will be only looking for convex piecewise linear RLFs in our study of biochemical networks. This means looking for vectors *c*_1_, …, *c_m_* ∈ ℝ^*ν*^ (for some positive integer *m*) such that 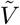 is an RLF where:

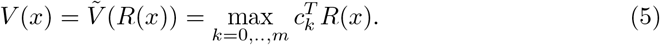

and *c*_0_ ≔ [0, ‥, 0]^*T*^. If the network has a positive steady state flux (i.e., there exists positive *r* such that Γ*r* = 0) then it can be shown that 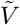 can be written as *V*(*x*) = ‖*CR*(*x*)‖_∞_, where ‖[*x*_1_, ‥, *x_n_*]^*T*^‖_∞_ ≔ max_*i*_ |*x_i_*| is the ∞-norm and 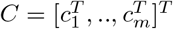. Two special cases are of interest to us:

#### Sum-of-Currents (SoC) RLFs

These are functions of the form:

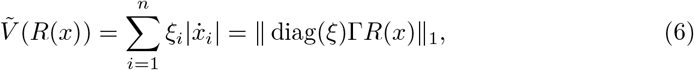

where 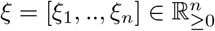 is a positive vector and ‖[*z*_1_, ‥, *z_n_*]^*T*^‖_1_ ≔ ∑_*i*_ |*z_i_*| is the 1-norm. The function considered in [22] is a special case with *ξ* = **1**. The vector *ξ* can be found by linear programming using a special case of Theorem 2 (see Methods). Note that the function (2) discussed in the motivating example has the form (6) above.

#### Max-Min RLFs

These are functions of the form:

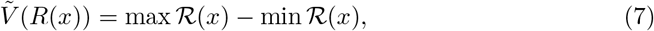

where 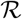 consists of reaction rates or the difference between forward and backward rates of a reaction. Unlike SoC RLFs which keep track of the reaction rate differences across each species, the Max-Min RLF keeps track of the maximal reaction rate difference across the *whole* network at each time. We provide a full graphical characterization of the class of networks that admit Max-Min RLFs (which we call *M*-networks). (see Methods, Theorem 4)

#### Alternative Forms

In Methods, we give conditions on a function 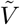 such that 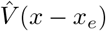 (where *x_e_* is a steady state) is a Lyapunov function for any admissible *R*. We call 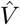 a concentration-dependent RLF. We show that 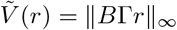 is an RLF iff 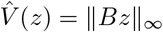 is a concentration-dependent RLF (see Methods, Theorem 11). These PWL functions relate to the ones proposed in [34, 36]. Note, however, that 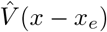 is a Lyapunov function only in the stoichiometric class that contains *x_e_*.

#### Properties of RLFs

In [27], some properties of networks admitting PWL RLFs have been established and they can serve as necessary condition tests. In Methods, we provide two additional properties, namely testing robust non-degeneracy and the absence of critical siphons. These conditions are implemented in `LEARN`.

### The Class of Structurally Attractive Biochemical Networks

The existence of an RLF implies that the qualitative long-term behavior of a network is highly constrained. Hence, an important issue is whether this theory is sufficiently relevant to biomolecular applications. We will show in the remainder of the Results section that this class of networks constitutes a rich and relevant class. It includes basic motifs, modules, and larger networks and cascades in molecular biology. For most of these networks, the HJF Lyapunov function [14] does not apply. And if it applies, it is only valid with Mass-Action kinetics (or a generalization [18]) and it does not confer the same powerful conclusions offered by our theory. Many of the networks discussed in the remainder of this paper are qualitatively analyzed for the first time and most of them had no Lyapunov functions known for them. For all the subsequent networks the following statement holds: if a positive steady state exists, then it is unique and globally asymptotically stable relative to its stoichiometric class.

### Binding/Unbinding Reactions

In this subsection, several biochemical networks are presented. They are fairly simple and all of them can be analyzed using HJF theory in the case of Mass-Action kinetics. However, they are presented here to show that the properties that our theory requires are obeyed by the basic biochemical motifs, which establishes its applicability and generality. Furthermore, we offer an intuitive window to the meaning of RLFs and how our graphical conditions apply.

#### Simple Binding Reaction

Figure 3-a represents a simple reversible binding reaction:

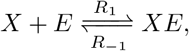

which can represent an enzyme binding to a substrate. The corresponding RLF can be found easily using Theorem 4 and is given by:

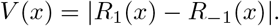

**Fig 3.**
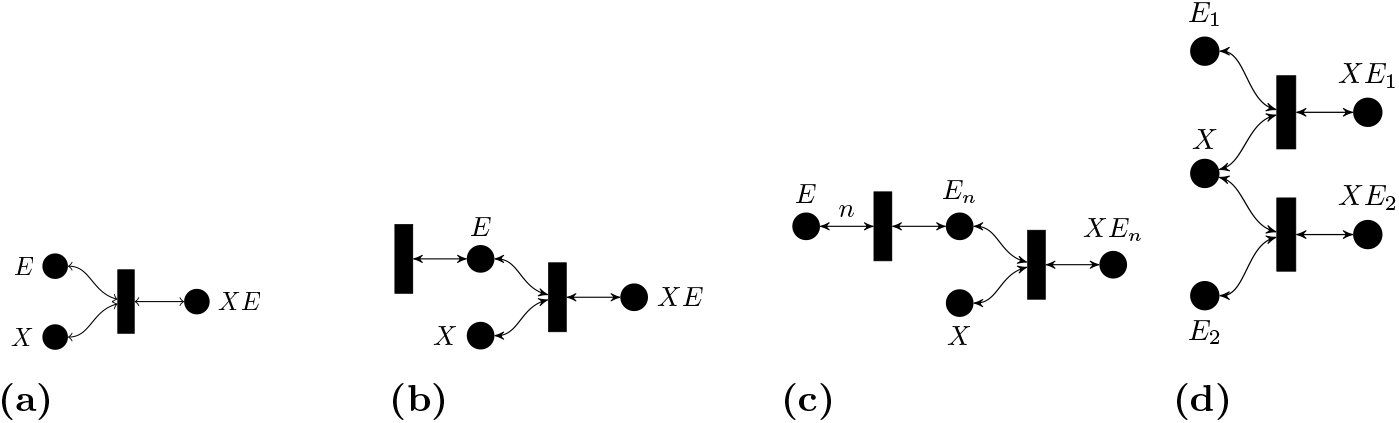
Basic Biochemical Examples. (a) Simple binding. (b) Simple binding with enzyme inflow-outflow. (c) Cooperative binding. (d) Competitive binding.

Both the Max-Min and the SoC RLFs coincide in this case.

#### Simple Binding With Enzyme Inflow-Outflow

Figure 3-b represents the following binding reaction with enzyme inflow-outflow:

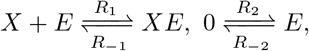

By considering the irreversible subnetwork 0 → *E*, 0 → *X, X* + *E → XE, XE* → 0, a Max-Min RLF can be found using Theorem 4 and is given by (7) where

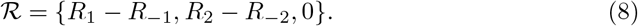

#### Cooperative Binding Reaction

The following reactions (depicted in Figure 3-c) represent the situation where *n* enzyme molecules *E* need to bind to each other to react to *X* :

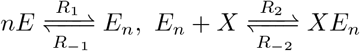

The case *n* = 2 is called dimerization. The corresponding RLF can be found using Theorem 4 and 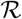 is given by (8). The irreversible subnetwork for which Theorem 4 was applied is 0 → *E*, 0 → *X, nE* → *E_n_*, *E_n_* + *X* → *XE_n_, XE_n_* → 0.

#### Competitive Binding Reaction

The following reactions (depicted in Figure 3-d) describe the situation when two molecules *E*_1_, *E*_2_ compete to bind with *X*:

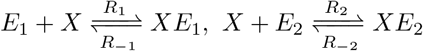

The corresponding RLF can be found using Theorem 4 and 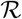 is given by (8). The irreversible subnetwork for which Theorem 4 was applied is 0 → *E*_1_, *E*_1_ + *X* → *XE*_1_ → *XE*_1_ → 0, 0 → *XE*_2_ → *X* + *E*_2_, *E*_2_ → 0.

### Three-Body Binding

We have applied our techniques to the dynamics of simple binding which can be analyzed easily using various known ways. However, it is often the case that two compounds *X, Y* cannot bind unless a bridging molecule *E* allows them to bind, forming a ternary complex. This is known as *three-body binding* [53] and it is ubiquitous in biology. Examples include T-cell receptors interaction with bacterial toxins [54], coagulation [55], and multi-enzyme supramolecular assembly [56]. The same reaction network also models the binding of two different transcription factors into a promoter with a double binding site. Despite its simplicity, the steady-state analysis of the equilibria has been subject of great interest [53]. Stability cannot be decided via HJF theory, and it has not been studied before to our knowledge.

The network can be depicted in Figure 4, and is given by eight reactions as follows:

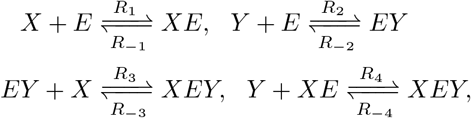

**Fig 4.**
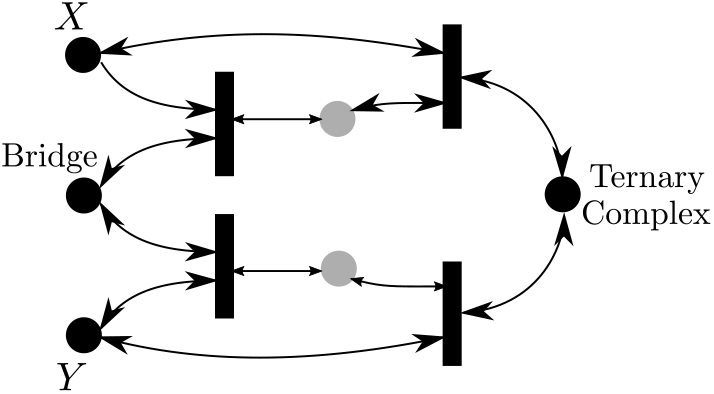
Three-Body Binding. Gray-colored species are intermediates.

The network is an *M*-network and the corresponding irreversible subnetwork has the reactions {**R**_1_, **R**_−2_, **R**_−3_, **R**_4_}. Hence we apply Theorem 4 to have an RLF of the form (7) where 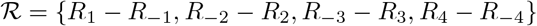. It can be concluded that there exists a unique steady state in each stoichiometric class and it is globally asymptotically stable.

### Transcription and Translation Networks

Transcription and translation are the first two essential steps in the central dogma of molecular biology, and hence they are of utmost importance in the analysis of gene regulatory networks.

#### Transcription Network

Figure 5-a) shows the transcription network which describes the production of mRNA from DNA using the RNA polymerase [57]:

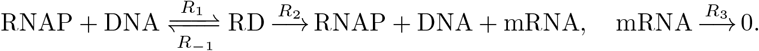

**Fig 5.**
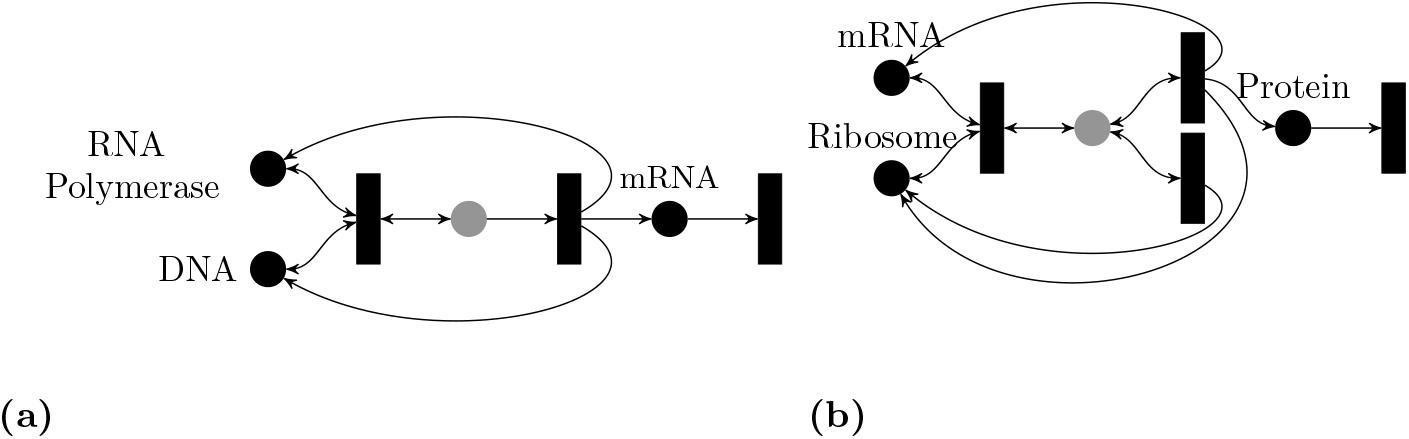
Transcription and Translation. Gray-colored species are intermediates. (a) Transcription. (b) Translation with a leak.

This model explicitly accounts for the concentration of RNA polymerase and hence it extends to situations in which RNA polymerase is not abundant.

Applying Theorem 4, the RLF (7) can be used with 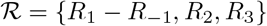. Alternatively, Theorem 2 can be used, and the Lyapunov function found can be written as: 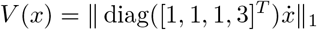, where the species are ordered as RNAP, DNA, RD, mRNA.

Note this network has deficiency one, hence no information regarding stability can be inferred from HJF theory. Furthermore, the procedure proposed in [6] has been reported not to work for the network above.

#### Translation Network With A Leak

Figure 5-b) shows the translation network which describes the production of a protein from mRNA via ribosomes [57]. The leaking of the Ribosome-mRNA complex into the pool of ribosomes is also modeled. In order to make the model more general, we also explicitly account for the concentrations of ribosomes. This is relevant to situations in which ribosomes are not highly abundant which can occur naturally [58, 59] or in synthetic circuits [60]. The network can be written as

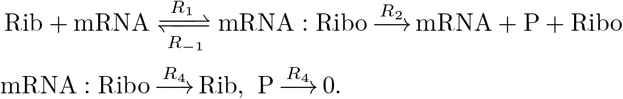

Note that the flux corresponding to reaction *R*_4_ vanishes at steady state which implies that the species mRNA:Ribo vanishes at any steady state. Note also that the dynamics of other species are independent of the dynamics of *P*. Hence, the network can be considered as a cascade of

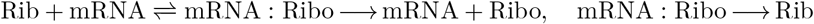

and 0 → *P* → 0. Applying Theorem 3 to the first network we get the following Lyapunov function:

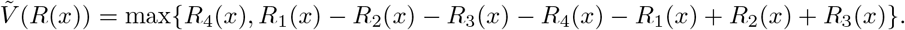

Note that 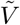 is neither SoC nor Max-Min. The second network can be analyzed using this Lyapunov function: *V*_2_(*x*) = |*R*_3_(*x*) − *R*_4_(*x*)|. Overall stability is established for the cascade using standard techniques [1].

### Basic Enzymatic Networks

#### Basic Activation Motif

Figure 6-a) represents the basic enzymatic reaction where an enzyme *E* binds to a substrate *S* to produce *S*^+^ as follows [48]:

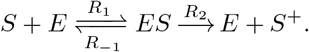

**Fig 6.**
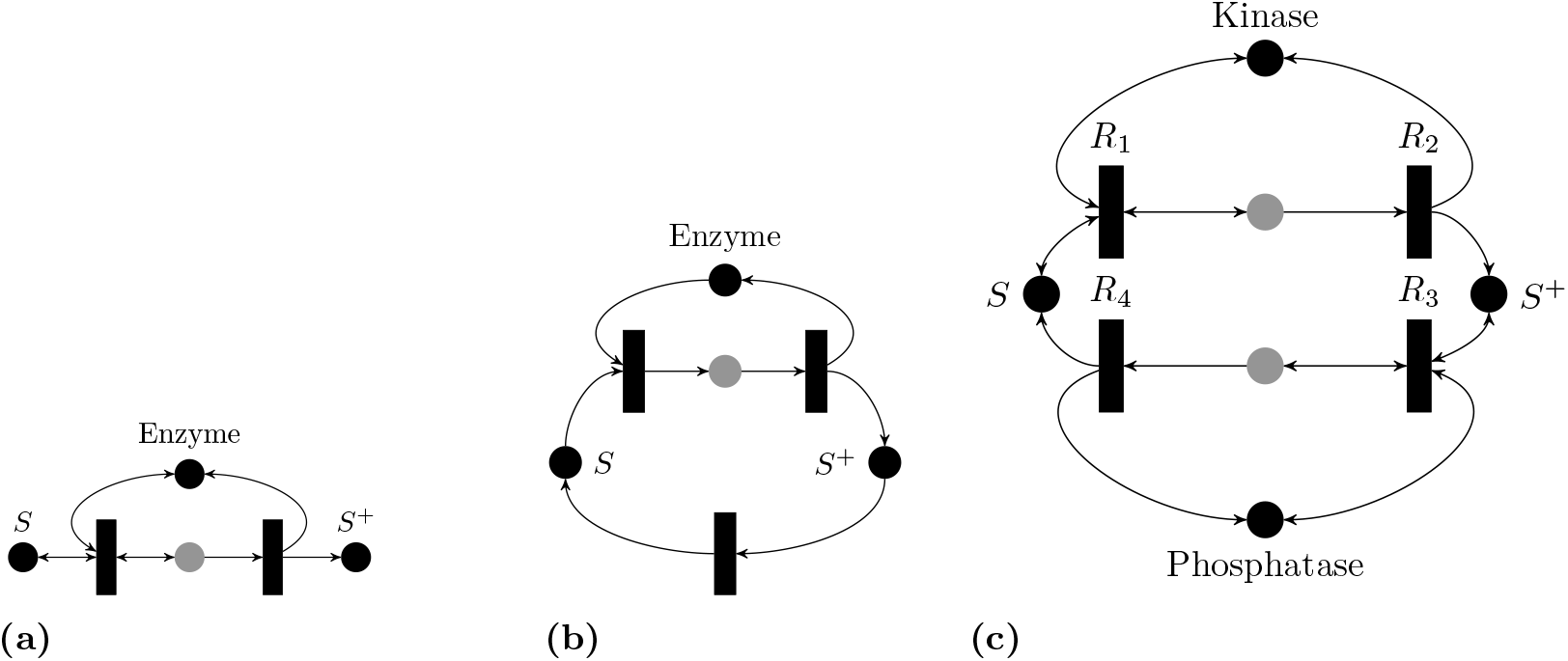
Basic Enzymatic Reactions. Gray colored species are intermediates. (a) Basic enzymatic motif. (b) Enzymatic cycle. (c) Full PTM cycle.

Theorem 3 can be used. The resulting Lyapunov function is: *V*(*x*) = max{|*R*_1_ − *R*_−1_|, *R*_2_}. Although this network has deficiency zero, it is not weakly reversible. This implies that the steady states belong to the boundary, and HJF theory does not offer any information regarding stability.

#### Enzymatic Activation Cycle

In order to close the cycle of the activation motif, Figure 6-c) depicts the activation of a protein *P* by an enzyme *E*, and then the activated protein decays back to its inactive state. The list of reactions is given as [2]:

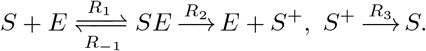

Theorem 2 gives the following SoC RLF:

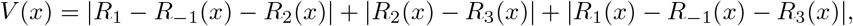

and both Theorems 3 and 4 give RLFs also.

This network has deficiency one; the deficiency one algorithm [17] excludes the existence of multiple steady states with Mass-Action kinetics. No information regarding stability can be inferred in that context from HJF theory. Furthermore, the decay reaction *R*_3_ usually models fast dephosphorylation which has a Michaelis-Menten kinetics, which is not allowed in [17].

#### The Full PTM Cycle

A simplified version of the enzymatic futile cycle has already been used as a motivating example in Figure 2. It differs from the preceding network by explicitly modeling the dephosphorylation step. The following describes the complete model [48, 49]:

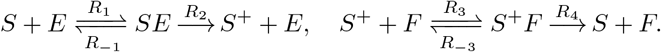

For instance, *S* represents the base substrate, *E* is called a kinase which adds a phosphate group to *S* to produce *S*^+^. This process is called *phosphorylation*. The *dephosphorylation* reaction is achieved by a phosphatase *F* that removes the phosphate group from *S*^+^ to produce *S*.

Theorem 4 can be used to find the RLF (7) where 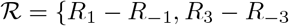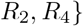.

Alternatively, Theorems 3 yields the SoC RLF:

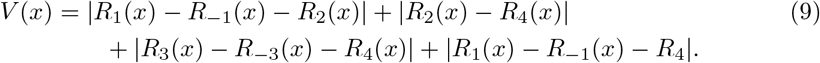

Both SoC and Max-Min RLFs have an intuitive meaning in terms of the reaction graphs of the networks. The first is the difference between the fastest and the slowest reactions, and the second is the sum of currents (rates of change of concentrations). Since the deficiency of the network is one, stability cannot be inferred from HJF theory.

#### Energy-constrained PTM Cycle

##### Basic Motif

Madhani [63] presents this biochemical example of adding a phosphate group to a protein using a kinase. ATP is not assumed to be abundant and its dynamics are explicitly modeled. The reaction network is depicted in black in Figure 7-a), which can be written as:

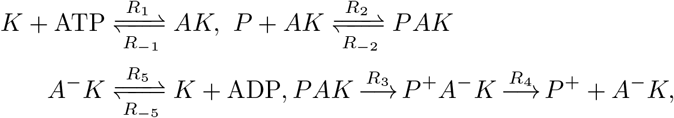

where *K* is the kinase, ATP is the adenosine triphosphate, ADP is the Adenosine diphosphate, and *P*^+^ is the phosphorylated protein. Reactions *R*_3_, *R*_4_ are not supported in the kernel of the stoichiometry matrix, which implies that the species *PAK, P* ^+^*A*^−^*K* vanish at any steady state point.

**Fig 7.**
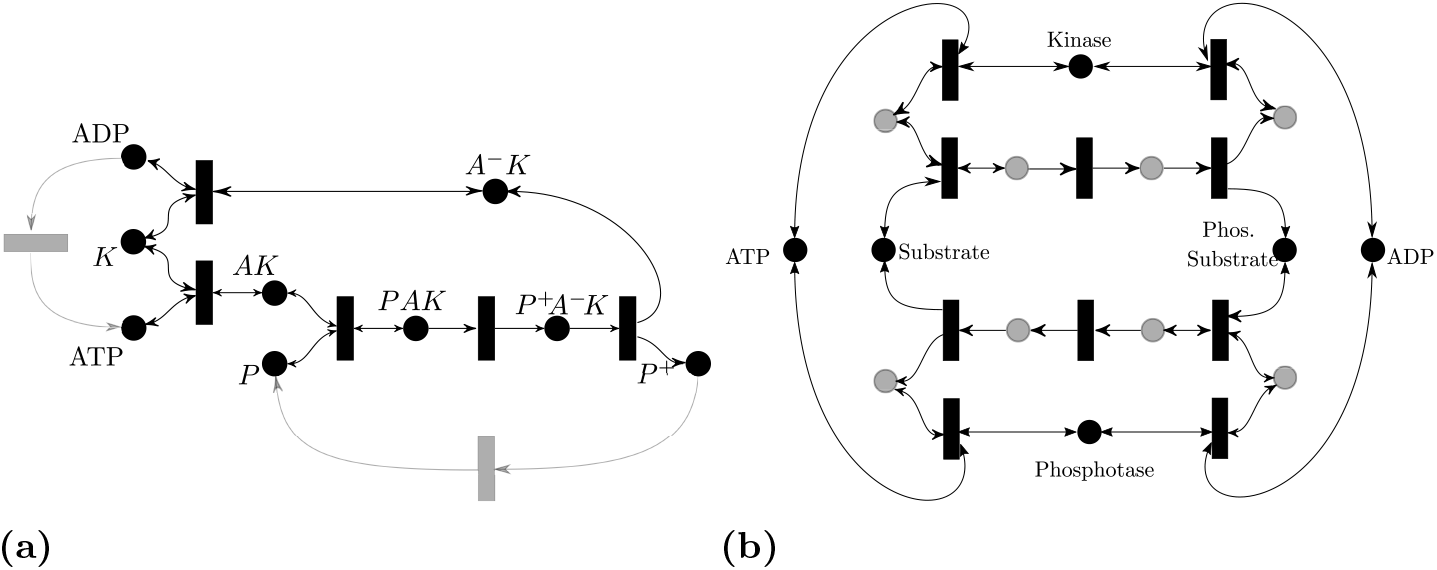
Energy-constrained PTM cycles. (a) Phosphorylation is modeled only. The black-colored component is the basic motif proposed in [63] (b) A full phosphorylation-dephosphorylation cycle with energy expenditures modeled. The gray species are intermediates.

Applying Theorem 3, one can get the following RLF function:

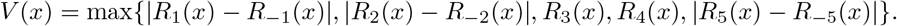

##### Energy Constrained PTM Cycle

In order to have a full cycle, the model can include the following two reactions: 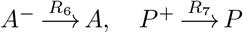, where ADP is converted to ATP by other cellular processes and is modeled as a single step, and *P*^+^ decays to its original state *P* spontaneously or chemically [64]. The reaction network is depicted in Figure 7-a).

The full network is an *M* network, and it has the RLF (7) with 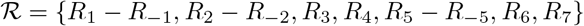.

The network is not complex-balanced and HJF theory is not applicable. The dynamics of this network have not been analyzed before per our knowledge.

##### Full Energy-Constrained PTM Cycle

The dephosphorylation step can be modeled fully and is depicted in Figure 7-b). This is the energy-constrained analog of Figure 6-c). The network is also an *M*-network and it admits an RLF of the form (7). The list of reactions have not been included for the sake of brevity.

### Post-Translational Modification Cycle Cascades

The post-translation modification (PTM) cycle (e.g, phosphorylation-dephosphorylation cycle [48, 49]) has been analyzed in the previous section. This kind of cycle appears frequently in biochemical networks, and can be interconnected in several ways; we discuss some here. For recent reviews see [65, 66].

#### A multisite PTM with Distinct Enzymes

It is known that a single protein can have up to different 100 different PTM sites [65] and it can undergo different PTM cycles such as phosphorylation, acetylation and methylation [67, 68]. Each of these cycles has its own enzymes.

Hence, we consider a cascade of *n* PTM cycles as shown in Figure 8-a) where *n* is any integer greater than zero. For instance, the associated reaction network for the case *n* = 2 is given as:

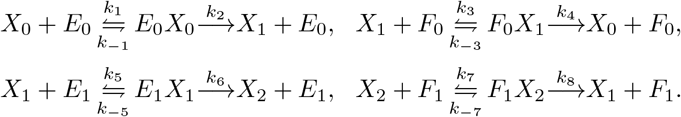

**Fig 8.**
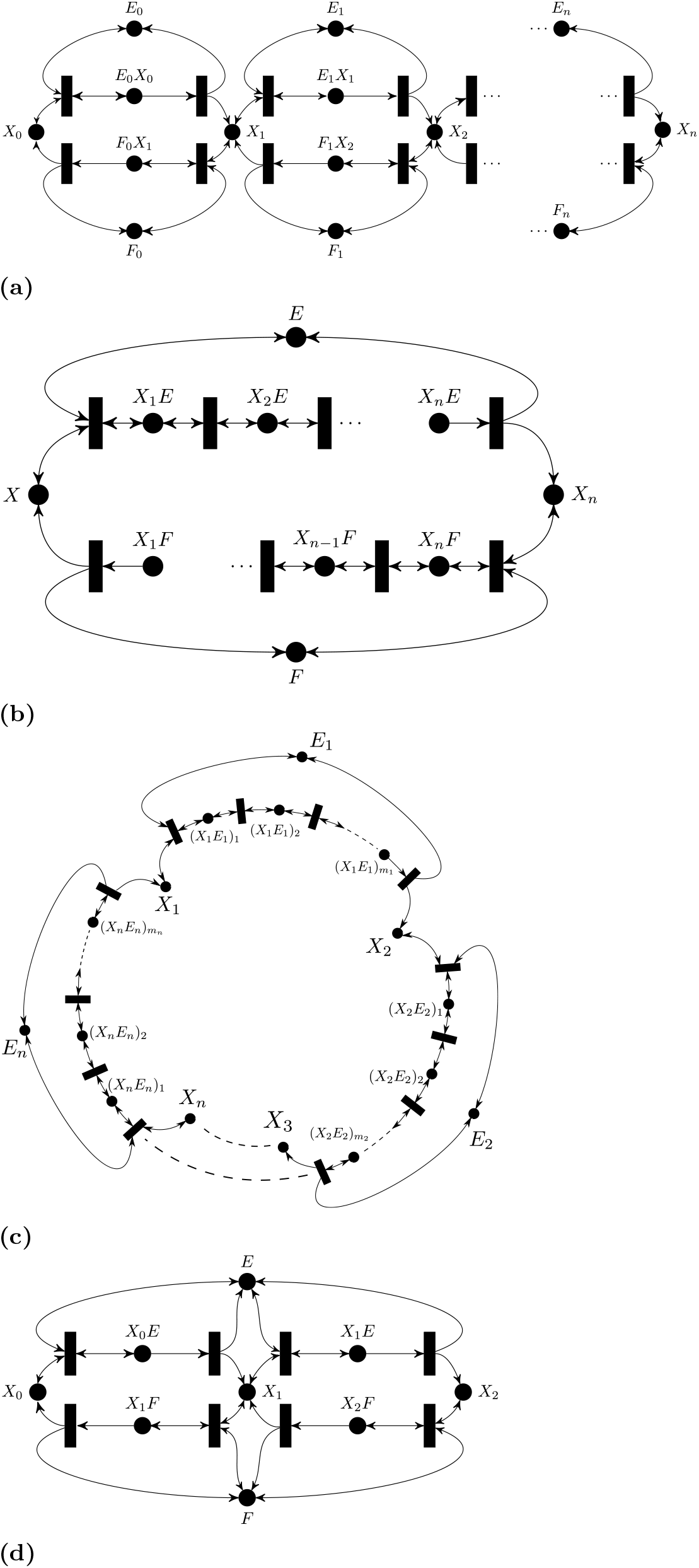
Cascades of PTM cycles. (a) A multisite PTM with distinct enzymes. (b) A multiple PTM with a processive mechanism. (c) The “all-encompassing” processive PTM mechanism. (d) Double PTM Cycle with a distributive mechanism.

The network is not an *M*-network and hence Theorem 4 is not applicable. However, using Theorem 2 it can be shown that a SoC RLF for the *n* cascade exists and can be represented as 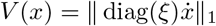 with *ξ* = [2, 2, …, 2, 1, 1, …, 1]^*T*^ with the ordering given as *X*_0_, …, *X_n_*, *E*_0_, *E*_1_, …, *F*_*n*−1_*X_n_*.

HJF theory will not apply since this network has deficiency *n*. Also, monotonicity-based results [24] do not apply, since the network is not cooperative in reaction coordinates. In fact, the long-term behavior of this cascade has not been studied before to our knowledge. It follows that for any *n* the network has a unique globally asymptotic stable steady state in any stoichiometric class (i.e, with respect to fixed total amounts for the enzymes and the substrate).

#### Multiple PTM cycles with a processive mechanism

Proteins can undergo different PTMs, but they also can undergo a multisite PTM. For instance, a phosphate group can be added to multiple sites on the protein [69]. Multisite phosphorylation can be processive [70] or distributive [71]. Figure 8-b) depicts a multiple-site futile cycle with a processive mechanism. The reaction network can be written as [33]

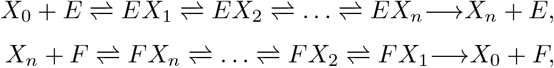

It can be noticed that for every *n*, the network satisfies the graphical conditions of Theorem 4. Therefore, an RLF is (7) where 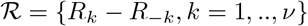, and 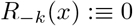 if **R**_*k*_ is irreversible.

##### Energy-Constrained Processive Cycle

The ATP and ADP expenditure can be accounted for in the processive cycle similar to the model presented in Figure 7-b). The new network will remain an *M*-network and Theorem 4 can be applied. Details are omitted for brevity.

##### A generalized processive cycle

An “all-encompassing” processive cycle has been studied in [8] which allows multiple enzymes and is depicted in Figure 8-c. It takes the following form:

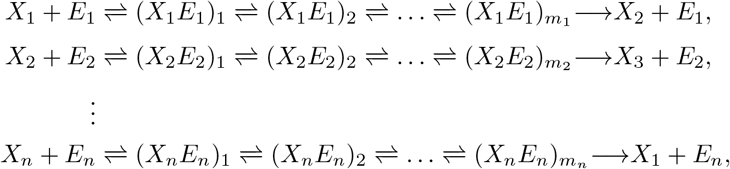

This network is also an *M* network and it satisfies the results of Theorem 4. Hence, the Lyapunov function (7) can be used.

Both networks above have been studied in [8, 33] by establishing monotonicity in reaction coordinates. Such techniques require checking persistence a priori and do not provide Lyapunov functions. Furthermore, our results have the advantage of providing an “all-encompassing” general framework that includes many of these individually studied networks in addition to new ones.

#### Distinguishing between Processive and Distributive Mechanisms

Figure 8-d) depicts a double futile cycle with a distributive mechanism [71, 72], which is described by the following set of reactions:

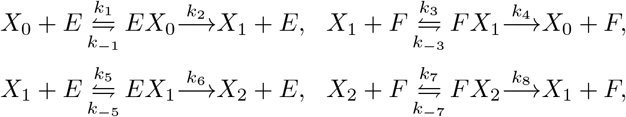

It can be verified that the network violates the *P*_0_ necessary condition (for the minor corresponding to *X*_0_, *X*_1_, *X*_2_, *E, FX*_1_, *EX*_1_). Hence, a PWL RLF does not exist [27]. Indeed, the above network is known to admit multi-stability for some parameter choices as shown in Figure 1.

Hence, our results can be used to compare between distributive and processive mechanisms as viable models for the first stage in the MAPK cascade. Since the latter has been observed experimentally to accommodate multiple non-degenerate steady states, the processive mechanism cannot be a model. (Similar observations have been made in [72–74].) Figure 1 depicts sample trajectories for the processive and distributive cycle with Mass-Action kinetics.

### Phosphotransfer and Phosphorelay Networks

Phosphotransfer is a covalent modification in which a histidine kinase gives the phosphate group to a response regulator and it is the core motif in a two-component signaling systems [75]. Phosphotransfer cascades are called phosphorelays [76, 77].

#### Phosphotransfer motif

An example is the envZ/ompR signaling system for regulating osmolarity in bacteria such as E. Coli [78]. The core motif can be described by the following set of reactions [79]:

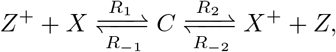

where the “+” superscript refers to a phosphorylated substrate. For instance, *Z*^+^ is the phosphorylated EnvZ protein, while *X* is the ompR protein.

The proteins *Z*, *X*^+^ can also be phosphorylated and dephosphorylated by other reactions. Figure 9-a) presents a network where those other reactions are modeled as a single step:

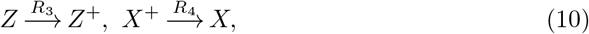

where *R*_3_ (which phosphorylates *Z*) can be monotonically dependent on external signals such as osmolarity in the envZ/OmpR network.

**Fig 9.**
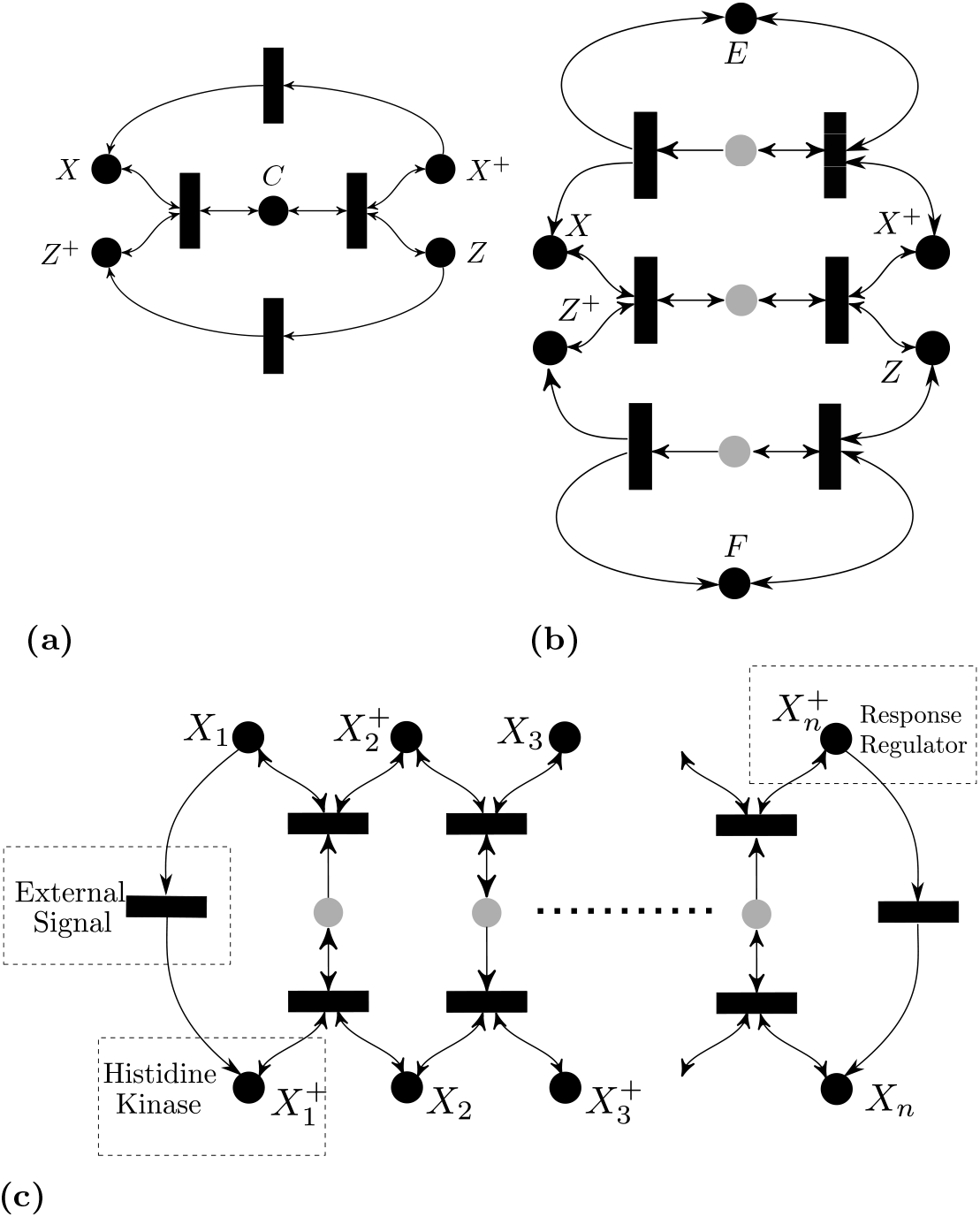
Phosphotransfer and Phosphorelay Networks. (a) Phosphotransfer network. (b) Phosphotransfer with phosphorylation/dephosphorylation. (c) A phosphorelay network.

It can be noticed that Theorem 4 is applicable and (7) is an RLF with 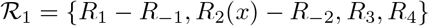.

#### Phosphotransfer with Enzymes

A more elaborate model can take into account the phosphorylation/dephosphorylation of proteins *Z, X*^+^ in terms of other enzymes. Hence, reactions (10) can be replaced by the following:

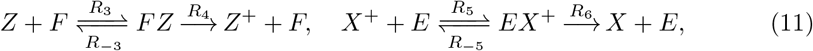

as depicted in Figure 9-b. Similarly, (7) is an RLF with 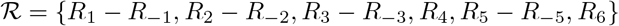.

#### A Phosphorelay

A phosphorelay is a cascade of several phosphotransfers. It appears ubiquitously in many organisms. For example, the KinA-Spo0F-Spo0B-Spo0A cascade in Bacillus subtilis [80] and the Sln1p-Ypd1p-Ssk1p cascade in yeast [81].

Figure 9-c depicts the cascade which is given by:

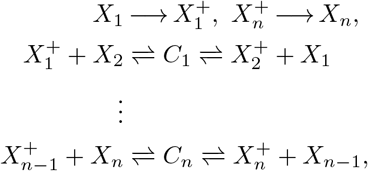

where the first kinase is phosphorylated by some constant external signal, and 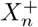 is the response regulator.

The network is still an *M*-network and conditions of Theorem 4 apply by mere inspection of the graph. Hence a function of the form (7) is a Lyapunov function. Enzymatic activation/deactivation of *X*_1_, 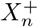, respectively, can also be added (analogously to Figure 9-b) and the result will continue to hold. Note that the same applies to the more general model presented in [82] also. We omitted the details for brevity.

Note that none of the phosphotransfer networks is complex-balanced and hence HJF theory is not applicable.

### T-Cell Kinetic Proofreading Network

In 1974, Hopfield [83] proposed the kinetic proofreading model in protein synthesis and DNA replication. Subsequently, McKeithan [84] proposed a network containing a ligand, which is a peptide-major histocompatibility complex *M*, binding to a *T*-cell receptor; the receptor-ligand complex undergoes several reactions to reach the final complex *C_N_*. The chain of reactions enhances the recognition and hence it is called a kinetic proofreading process. Figure 10-a) depicts the reaction network, which is given by the following set of reactions:

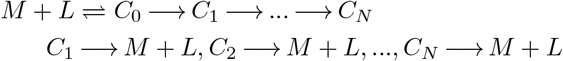

**Fig 10.**
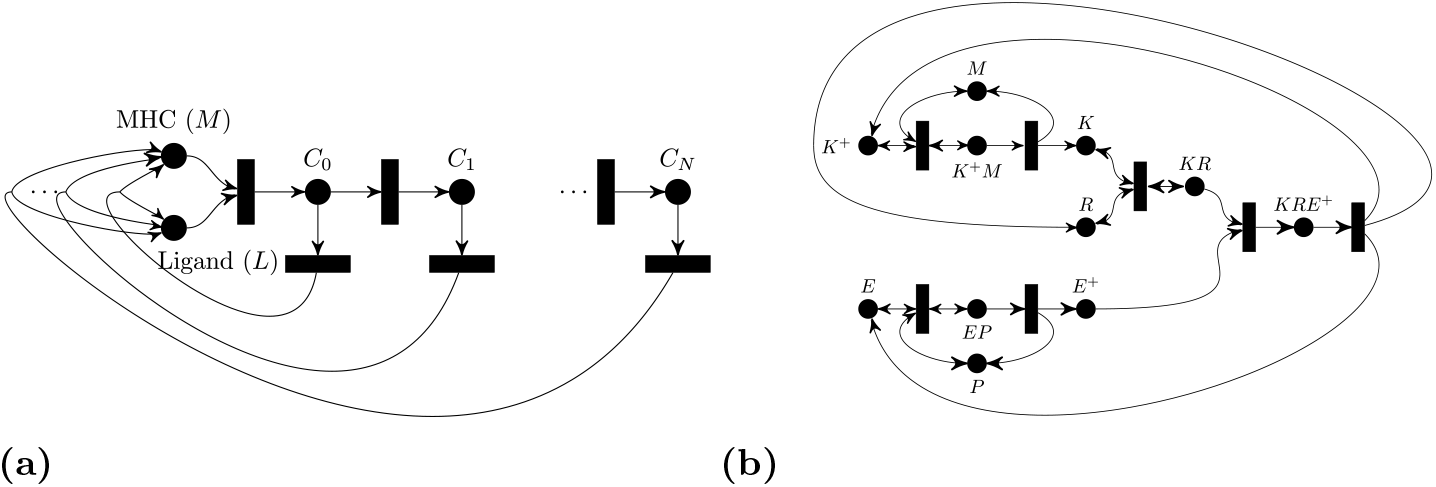
Other signalling networks. (a) McKeithan’s T-Cell kinetic proofreading network. (b)ERK signaling Pathway With RKIP Regulation

Applying Theorem 2, it can be shown that for any *N* ≥ 1, the network admits a SoC RLF of the form 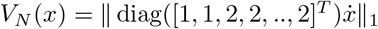, where the species are ordered as *T, L, C*_0_, *C*_1_, …, *C_N_*. Note that this network does not meet the graphical requirements of Theorem 4 since it is not an *M* network. The monotone-systems approach proposed in [24] is not applicable here since the system is not cooperative in reaction coordinates.

Nevertheless, this is one of the few networks, considered so far, which is complex-balanced. The work [18] showed that this network is weakly reversible and that it has zero-deficiency; therefore any positive steady state is unique relative to the interior and is locally asymptotically stable. In order to infer global stability, it was necessary to compute the steady states explicitly to preclude a boundary steady state stoichiometrically compatible with a positive steady state. In comparison, our approach is more powerful, since the former approach is limited to generalized Mass-Action kinetics, and cannot infer global stability directly.

### ERK signaling Pathway with RKIP Regulation

Figure 10-b depicts the network describing the effect of the so called Raf Kinase Inhibitor Protein (RKIP) on the Extracellular Regulated Kinase (ERK) signaling pathway as per the model given in [85]. It can be described using the network:

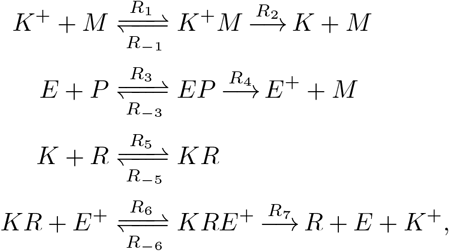

where *K* is the RKIP, *E* is the ERK Kinase, *P* is the RKIP phosphatase, and *M* is the phosphorylated MAPK/ERK Kinase, and the plus superscript means that the molecule is phosphorylated.

The network is an *M*-network and the requirements of Theorem 4 are satisfied. Hence, (7) is an RLF with 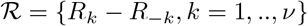, and *R*_−*k*_(*x*) : 0 if **R**_*k*_ is irreversible. Note that this network is of deficiency one, hence stability cannot be inferred by HJF theory. Nevertheless, monotonicity-based analysis can be applied [4] which utilizes cooperativity in reaction coordinates. Refer to the Discussion for a detailed comparison to monotonicity techniques.

### The Ribosome Flow Model

Finally, we show that our techniques’ applications in molecular biology are not limited to classical biochemical networks. A translation elongation process involves ribosomes travelling down an mRNA, readings codons and translating amino-acid chains via recruited tRNAs. A conventional stochastic model is the *Totally Asymmetric Simple Exclusion Process* [86]. A coarse-grained mean-field approximation that resulted in a deterministic continuous-time flow model was introduced by [87], and its dynamics have been studied further [87, 88].

Figure 11 illustrates the model. An mRNA consists of codons that are grouped into *n* sites, each site *i* has an associated occupancy level *x_i_*(*t*) ∈ [0, 1] which can be interpreted as the probability that the site is occupied at time *t*. The ribosomes’ inflow to the first site is *λ*_0_, which is known as the initiation rate, *λ_i_* is the elongation rate from site *i* to site *i* + 1, and *λ_n_* is the production rate. All rates are assumed to be positive. The ODE is written as follows:

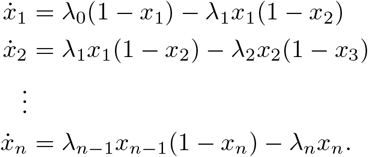

**Fig 11.**
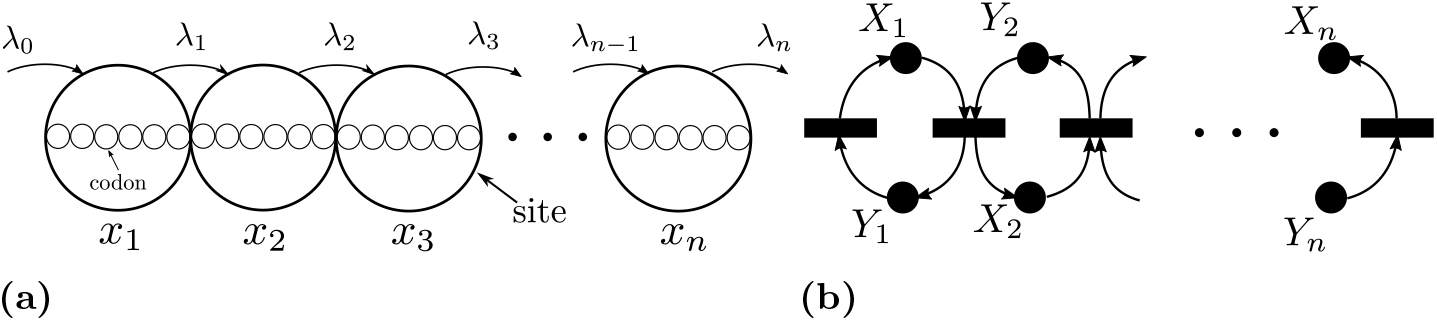
The Ribosome flow model. (a) Schematic representation. *λ*_0_ is the initiation rate, *λ_i_* is the elongation rate from site *i* to site *i* + 1, and *λ_n_* is the production rate. The state variable *x_i_* ∈ [0, 1] is the occupancy level of the site *i*. (b) Reaction network representation. *X_i_* corresponds to the occupancy level, while *Y_i_* corresponds to the vacancy level.

The dynamics of the system above have been analyzed and shown to be monotone in [88]. In what follows, we provide an alternative approach that provides a Lyapunov function and establishes more powerful properties. Let *y_i_* ≔ 1 − *x_i_*, *i* = 1, ‥, *n*. Then, we can define a reaction network with species *X_i_*, *Y_i_*, *i* = 1, ‥, *n* as follows:

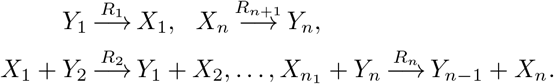

The network has 2*n* species, *n* + 1 reactions, and *n* conservation laws. It is depicted in Figure 11-(b). The ODE system above describes the time-evolution of the reaction network with Mass-Action kinetics.

The graphical conditions of Theorem 4 are satisfied. Hence, (7) is an RLF for any *n* with 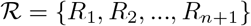. Since the network is conservative it follows that there exists a unique globally asymptotically stable steady state. Note that this results holds with general monotone kinetics.

### Quantitative analysis via RLFs

In this subsection we show that our RLFs can provide valuable quantitative information regarding the behavior of the network beyond mere qualitative long-term behavior information.

#### Safety sets

Since our techniques are based on the construction of RLFs, we can compute safety sets which are the level sets of a Lyapunov function. If a system starts in a safety set it cannot leave it at any future time. Substituting Mass-Action kinetics, the safety set for a Lyapunov function 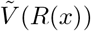 consists of piecewise polynomial surfaces and it is not necessarily convex. The safety set provided by an RLF surrounds all the steady states, i.e is not restricted to stoichiometric classes. In comparison, a concentration-dependent RLF provides a convex polyhedral safety set in a specific stoichiometric class. In order to illustrate this, consider the full PTM cycle with Mass-Action kinetics and let all the kinetic constants be 1. There are three conserved quantities, which we assume are set to [*E*]_*T*_ = [*F*]_*T*_ = [*S*]_*T*_ = 10 AU. Hence, the dynamics of the ODE evolve in a subset of three dimensional cube [0, 10]^3^. A level set of the RLF in (9) can be calculated restricted to the stoichiometric compatibility class and is depicted in the Figure 12-a. The concentration-dependent RLF can be constructed via Theorem 11. Plotting the level set requires computing the steady state which can be calculated by solving the algebraic equations to be: (*x_e_*, *e_e_*, *f_e_*) ≈ (1.216990, 6.216990, 6.216990). The level set is plotted in Figure 12-b. Both safety sets corresponding to the two Lyapunov functions are chosen so that *s* = 2.5 lies on the boundary of the set. In other words, the substrate concentration is guaranteed not to exceed 2.5 if the system is initialized in the set. It can be clearly seen that the two sets are distinct, and they give different guarantees. Their intersection gives a tighter safety set.

**Fig 12.**
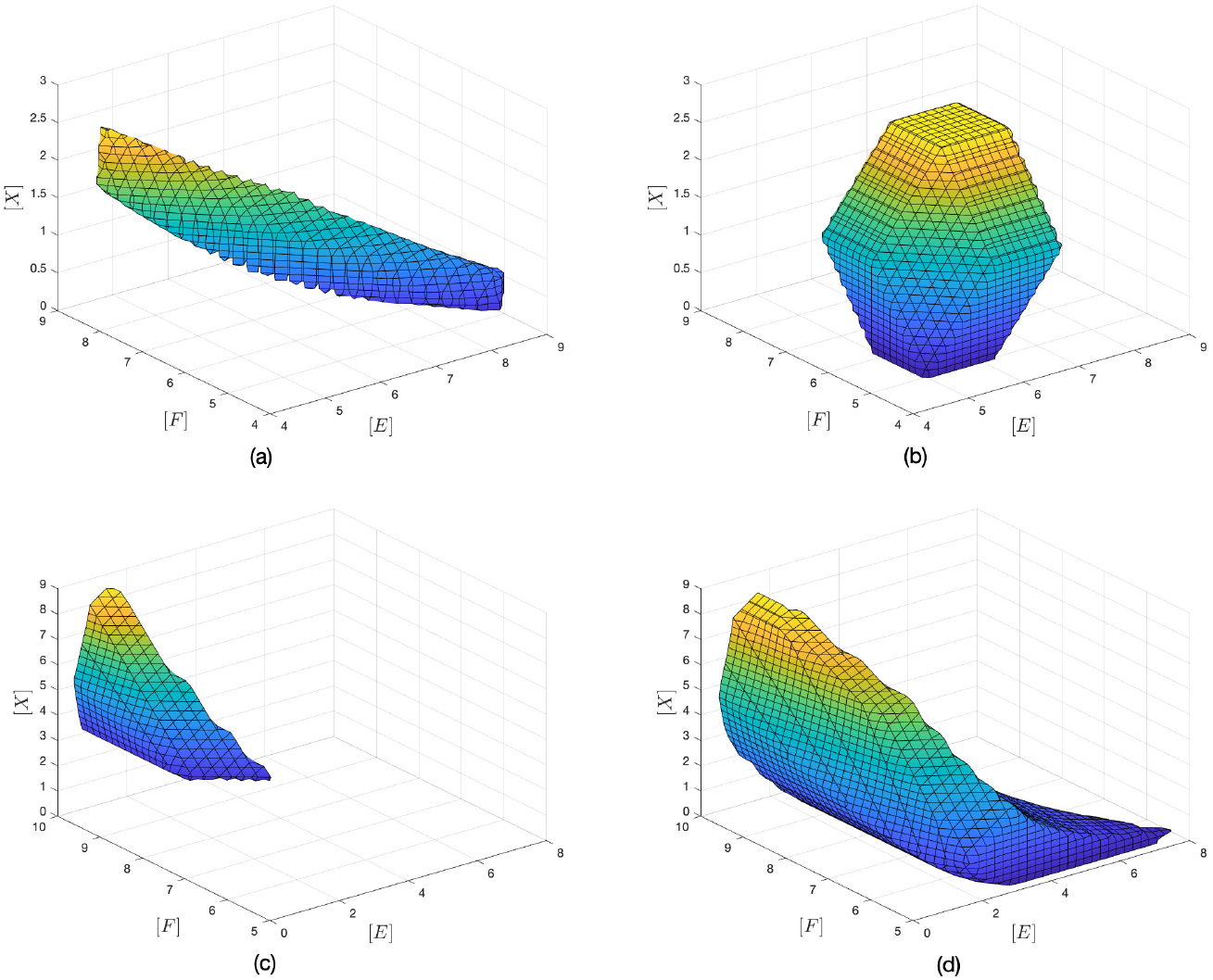
Safety sets computed via RLFs. **(a),(b)**, Safety sets for the PTM cycle (Figure 6-c). (a) The safety set corresponding to the rate-dependent RLF for the PTM cycle. It is the *α*-level set of *V* where *α* has been chosen such that the concentration of *S* does not exceed 2.5. (b) The safety set corresponding to the concentration-dependent RLF. The safety set has been chosen similarly to satisfy the same condition. **(c),(d)**, Sub-levels sets for the safety sets corresponding to the rate-dependent RLF (7) for the double processive PTM cycle (Figure 8-b). (c) The sublevel set (with [*X*_2_] = 0) of the *α*-level set of *V* where *α* has been chosen such that the concentration of *E* does not exceed 2.5 on the sublevel set. (d) Another sublevel set of the same set in (c) with [*X*_2_] = 0.5AU.

Another example is a double processive PTM (Figure 8-b) which has four dimensional stoichiometric classes. Hence, the 4D safety sets cannot be plotted, but their sublevel sets can still be visualized. Figure 12-c,d shows *sublevel* sets for different concentrations for the double phosphorylated species *X*_2_ with total kinase, phosphatase and substrate concentrations fixed to 10AU each. Figure 12-c) shows the safety set with the concentration of the free kinase *E* not exceeding 2.5 and with [*X*_2_] = 0. However, the sublevel set changes drastically if the concentration of *X*_2_ is 0.5AU as shown in Figure 12-d.

#### Flux analysis for the McKeithan Network

Since the RLF are written in terms of rates (also called fluxes), our functions can be used in the context of flux analysis. Such techniques usually operate at steady state and do not take dynamics into consideration [89]. We provide an illustrative example to show how our RLF can be used. Let *N* = 2 for the network above. Usually, the network is initialized with zero concentration of the intermediate complexes. Hence, the initial concentrations of *M, L* are [*M*]_*T*_, [*L*]_*T*_. Therefore, the Lyapunov function provides the following safety set 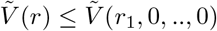, where *r*_1_ is the flux which is a function of [*M*]_*T*_, [*L*]_*T*_. For each [*M*]_*T*_, [*L*]_*T*_, we want to find an upper bound that *c*_2_ cannot exceed for all time. Let **R**_6_ be the last reaction (i.e, *C*_2_ → *M* + *L*), and let **R**_1_ be the first reaction, i.e *M* + *L* → *C*_0_. Hence, we look for solving the following convex optimization problem for a given 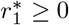:

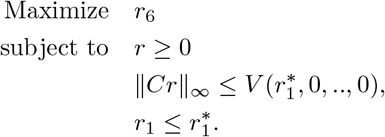

The last inequality is included since the network is conservative and *R*_6_(*m, ℓ*) ≤ *R*_6_([*M*]_*T*_, [*L*]_*T*_) holds due to the monotonicity of *R*.

The optimization problem above does not require knowledge of the kinetics as it is defined for fluxes. For the *T*-cell network, the solution of the problem is 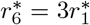. This means that the flux *r*_6_ is guaranteed to be less than 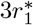 for all time. Converting these bounds to concentrations requires usage of the kinetics. Let *R*_1_(*m, ℓ*) = *k*_1_*mℓ*, and let 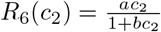 (Michaelis-Menten kinetics). Solving for *c*_2_, we can plot an upper bound on total amount of *k*_1_[*M*]_*T*_ [*L*]_*T*_ versus the maximum allowed concentration *c*_2_. If *R*_6_ is Mass-Action then the relationship will be linear. Both curves are plotted in Figure 13.

**Fig 13.**
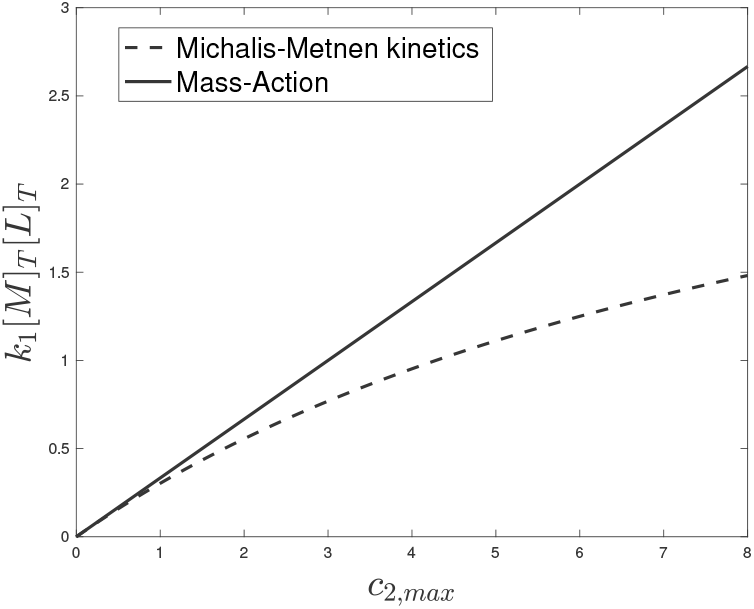
Flux analysis for McKeithan’s T-Cell kinetic proofreading network. The plot depicts an upper bound on the input flux versus the maximum allowed concentration of the end product with Michaelis-Menten kinetics *R*_6_(*c*_2_) = *c*_2_/(0.1*c*_2_ + 1) and Mass-Action kinetics *R*_6_(*c*_2_) = *c*_2_.

## Discussion

We have presented a comprehensive theoretical framework and provided computational tools for the identification of a class of “structurally attractive” networks. It has been demonstrated that this class is ubiquitous in systems biology. Networks in this class have universal energy-like functions called Robust Lyapunov Functions and, under additional mild conditions, can only admit unique globally stable steady states. Their Jacobians are well behaved and they cannot exhibit chaos, oscillations or multistability. The latter cannot be admitted even under inflow/outflow perturbations. Hence, LEARN can be used to rule out these networks as viable models for mechanisms that display such behaviors experimentally. Thus, our work supplements other mathematical methods used to invalidate models, as for example those in [90] and [91].

Our class of networks is distinct from the one identified by the HJF theory [14, 17] and it has wider applications to biology as we have shown. Furthermore, our results include all networks that have been studied via compartmental system techniques [22, 23] and via monotonicity techniques [8, 24, 33]. In fact, showing that the latter class of network always admits an RLF is a subject of a forthcoming paper. Refer to Table 1 for a comparison with techniques in the literature. In addition to wider applicability, our analysis has the advantage of showing persistence automatically, rather than needing to check it a priori as in [24]. Also, it has the advantage of having an explicit expression for the Lyapunov function which can be used for a deeper study of the dynamics such as the construction of safety sets and flux analysis as discussed before. In addition, Lyapunov functions have been extensively used to study the effect of interconnections, uncertainties, disturbances, and delays [9, 10].

Our study of biochemical networks is not meant to be exhaustive, since we only focused on common motifs and cascades. We provide a computational package to help the wider community apply our techniques to study new networks.

We have presented the RLFs with two representations: rate- and concentration-dependent, and we have provided a toy example for dynamic flux analysis via a rate-dependent RLF. We look forward to these results being developed further to complement standard flux analysis techniques.

For a given network, we have presented sufficient conditions for the existence of an RLF, and several necessary conditions. However, there are important networks that lie in the gap between the necessary and sufficient conditions. A relevant example is a ligand (L) binding a receptor (R), and initiating a PTM cycle for a substrate (S). The reaction network is:

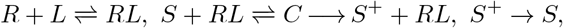

It satisfies all necessary conditions but its global stability is still open.

Future work includes the development of more general techniques to identify classes of networks that can be multi-stable but cannot admit oscillations or chaos. Furthermore, networks that admit RLFs have other strong properties in terms of contraction and stabilization [92], which will be studied in forthcoming papers.

## Methods

### Reaction Networks

We follow the standard notation and terminology on reaction networks [17, 18, 52, 93].

A Chemical Reaction Network (CRN) consists of *species* and *reactions*. A species is what participates or is produced in a chemical interaction. In the context of biochemical networks a species can be a gene’s promoter configuration, a substrate, an intermediate complex, an enzyme, etc. We denote the set of species by 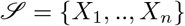. A reaction is the transformation of reactants into products. Examples include binding/unbinding, decay, complex formation, etc. We denote the set of reactions by 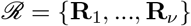. Reactions have two distinct elements: *the stoichiometry* and *the kinetics*.

#### Stoichiometry

The relative number of molecules of reactants and products between the sides of each reaction is the *stoichiometry*. Hence, each reaction is customarily written as follows:

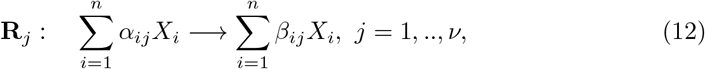

where *α_ij_*, *β_ij_* are nonnegative integers called *stoichiometry coefficients*. The expression on the left-hand side is called the *reactant complex*, while the one on the right-hand side is called the *product complex*. If a transformation is allowed to occur also in the opposite direction, the reaction is said to be *reversible* and its reverse is listed as a separate reaction. For convenience, the reverse reaction of **R**_*j*_ is denoted as **R**_−*j*_. The reactant or the product complex can be empty, though not simultaneously. An empty complex is denoted by 0. This is used to model external inflows and outflows.

An autocatalytic reaction is one which has a species appearing on both sides of the reaction simultaneously (e.g, *D → D* + *M*). A network is called *non-autocatalytic* if it has no autocatalytic reactions.

The stoichiometry of a network can be summarized by arranging the coefficients in an augmented matrix *n* × 2*ν* as: 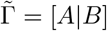, where [*A*]_*ij*_ = *α_ij_*, [*B*]_*ij*_ = *β_ij_*. The two submatrices *A, B* can be subtracted to yield an *n* × *ν* matrix 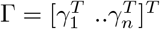 called the *stoichiometry matrix*, which is defined as Γ = *B* − *A*, or element-wise as: [Γ]_*ij*_ = *β_ij_* − *α_ij_*.

#### Kinetics

The relations that determine the velocity of transformation of reactants into products are known as *kinetics*. We assume an isothermal well-stirred reaction medium. In order to study kinetics, a nonnegative number *x_i_* is associated to each species *X_i_* to denote its *concentration*. Assume that the chemical reaction **R**_*j*_ takes place continuously in time. A *reaction rate* or velocity function 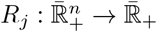 is assigned to each reaction. The widely-used Mass-Action kinetics have the following expression: 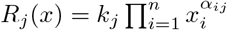, where *k_j_*, *j* = 1, ‥, *ν* are positive numbers known as the *kinetic constants*. Many other kinetic forms are used in biology such as Michaelis-Menten, Hill kinetics, etc.

We do not assume particular kinetics. We only assume that the reaction rate functions *R_j_*(*x*), *j* = 1, ‥*ν* satisfy the following minimal assumptions:

**AK1.** each reaction varies smoothly with respects to its reactants, i.e *R_j_*(*x*) is continuously differentiable;
**AK2.** each reaction needs all its reactants to occur, i.e., if *α_ij_* > 0, then *x_i_* = 0 implies *R_j_*(*x*) = 0;
**AK3.** each reaction rate is monotone with respect to its reactants, i.e *∂R_j_*/*∂x_i_*(*x*) ≥ 0 if *α_ij_* > 0 and *∂R_j_*/*∂x_i_*(*x*) ≡ 0 if *α_ij_* = 0;
**AK4.** The inequality in AK3 holds strictly for all positive concentrations, i.e when 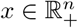.

Reaction rate functions satisfying AK1-AK4 are called *admissible*. For given stoichiometric matrices *A, B*, the set of admissible reactions is denoted by 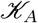.

#### Dynamics

The dynamics have been already given in (1). The set 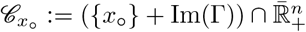 is forward invariant for any initial condition *x*_◦_, and it is called *the stoichiometric compatibility class* associated with *x*_◦_. For a conservative network all stoichiometric classes are compact convex polyhedral sets.

We sometimes will use the following assumption which is necessary for the existence of *positive* steady states.

**AS1** There exists *v* ∈ ker Γ such that *v* ≫ 0.

### RLFs and the Decomposition of the Dynamics

We have provided an informal definition of the notion of RLF in the introduction. The inequality in Eq. (4) must hold for all 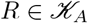. As observed before, AK1-AK4 imply a zero-sign pattern on *∂R/∂x* (see Figure 2-d for an illustration). This motivates defining the class of matrices with the specific sign pattern as follows:

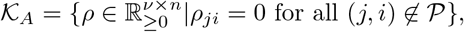

where 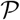 is the set of reaction-reactant pairs defined before.

#### Definition 1.

*Given a network* 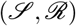. *A locally Lipschitz function* 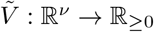 *is said to be an RLF if it satisfies the following:*

1. 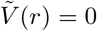 *iff r* ∈ ker Γ.
2. 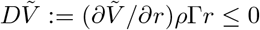 *for all* 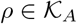 *and all r for which* 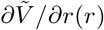 *exists*.

At points of non-differentiability, the time-derivative of 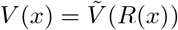 is defined in the sense of Dini (see S1 Text §1.1 for a review of Lyapunov theory and generalized derivatives).

We will show how the rank-one matrices *Q*_1_, ‥, *Q_s_* (defined in the Results section) can be used to embed the dynamics of the nonlinear network in a cone of linear systems. Although the Lyapunov function 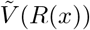 is a function in the concentration *x*, it is defined as a composition 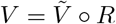. Therefore, we study the ODE in reaction coordinates. Let *x*(*t*) be a trajectory that satisfies (1) and let *r*(*t*) ≔ *R*(*x*(*t*)). Hence,

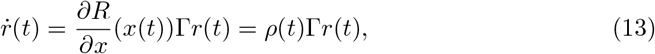

where 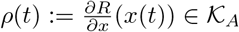.

The basic idea is to consider *ρ*(*t*) as an unknown time-varying matrix. Since its zero-sign pattern is known, we can decompose *ρ*(*t*) in the following way:

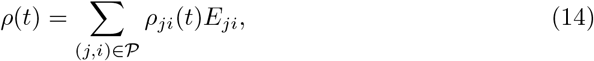

where [*ρ*(*t*)]_*ji*_ = *ρ_ji_*(*t*) > 0, and 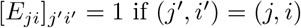 and zero otherwise. The matrices {*E_ji_*|(*i, j*) such that *α_ij_* = 0} form the canonical basis of the matrix space 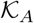.

Substituting (14) in (13) we can embed the dynamics of the network (13) in the conic combinations of a finite set of extremal linear systems as follows:

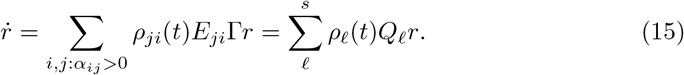

where *Q*_*ℓ*_, *ℓ* = 1, ‥, *s* have been defined before Theorem 1. This also implies that the Jacobian of (13) can be written at any interior point as: 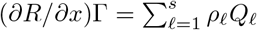. Hence, the Jacobian belongs to cone generated by the extremals *Q*_1_, ‥, *Q_s_*. Note that (15) can be interpreted as representing a linear parameter-varying system which has *s* nonnegative time-varying parameters {*ρ*_1_(*t*), ‥, *ρ_s_*(*t*)}. The linear systems are given by rank-one extremals *Q*_1_, ‥, *Q_s_*. The proof of Theorem 1 is completed in S1 Text §1.2.

### Computational Construction of RLFs

The results presented in [26, 27] have been derived via a direct analysis of the associated reaction networks. The framework introduced above enables interpreting these results in a more general framework and allows generalizing them. Hence we revisit the algorithms introduced for the existence and construction of PWL RLFs, and implement them in the `LEARN` MATLAB package. Furthermore, we also introduce piecewise quadratic RLFs based on the new framework introduced in this paper.

#### Piecewise Linear RLFs

Consider a CRN (1) with a Γ ∈ ℝ^*n*×*r*^ and a given *partitioning* matrix *H* ∈ ℝ^*p*×*r*^ such that ker *H* = ker Γ. A PWL RLF is piecewise linear-in-rates, i.e., it has the form: 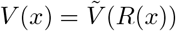, where 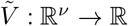 is a continuous PWL function. Assuming AS1, the piecewise linear function is given as

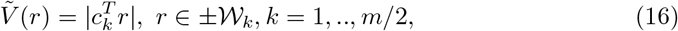

where the regions 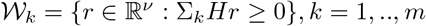 form a proper conic partition of ℝ^*ν*^, while 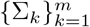 are signature matrices (diagonal matrices with ±1 on the diagonal) with the property Σ_*k*_ = −Σ_*m*+1−*k*_, *k* = 1, ‥, *m*/2. The coefficient vectors of each linear component can be collected in a matrix 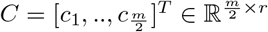. If the function 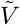 is convex, then we have the following simplified representation of *V* :

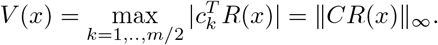

##### Verifying a candidate RLF

Checking if a given PWL function is an RLF can be posed as a linear program. It is discussed in S1 Text §2.1 and is coded into `LEARN`.

#### Construction via Linear Programming

Based on Theorem 1, we present a simpler linear program than the one presented in [27]. The proof is presented in S1 Text §2.2.

##### Theorem 2.

*Given a network* 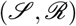 *that satisfies AS1 and a partitioning matrix H* ∈ ℝ^*p*×*r*^. *Let* {*v_i_*} *be a basis for* ker Γ. *Consider the linear program:*

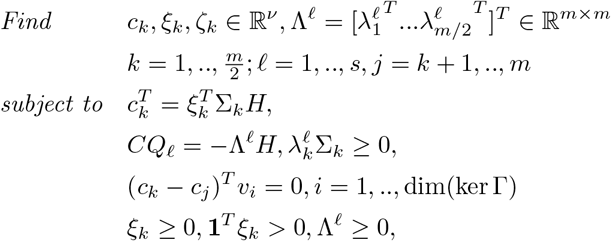

*Then there exists a PWL RLF with partitioning matrix H if and only if there exists a feasible solution to the above linear program that satisfies* ker *C* = ker Γ.

##### Remark 1.

*The linear program above does not enforce convexity on* 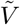. *Nevertheless, `LEARN` allows the user to search amongst convex* 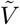*’s only. See S1 Text §2.3*.

In `LEARN` there is a default choice for the matrix *H*, and it also allows for a manual input by the user. The default choice is *H* = Γ which gives the following Lyapunov function (where the SoC RLF introduced in (6) is a special case):

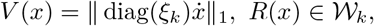

The user can add rows to *H*. Usually rows of the form {*γ_i_* ± *γ_j_*|*i, j* = 1, ‥, *n*}, where *γ*_1_, ‥, *γ_n_* are the rows of Γ, are good candidates.

##### Networks without positive steady states

If AS1 is not satisfied, then a linear program can be designed for constructing RLFs over a given partition. This is discussed in S1 Text §2.4.

#### An iteration for the construction of convex PWL RLFs

Assuming both AS1 and allowing non-autocatalytic networks only, a computationally-light iterative algorithm for constructing a convex Lyapunov function was presented in [26, 27]. Here we generalize the algorithm by dropping these two assumptions. The objective is to find a matrix 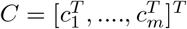 such that 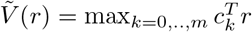 is a Lyapunov function, where *c*_0_ ≔ **0**.

We state the algorithm below. We use the notation supp(*c_k_*) = {**R**_*j*_|*c_kj_* ≠ 0}, which is the set of all those reactions that appear in 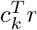, and let 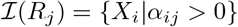 which is the set of reactants for reaction **R**_*j*_. We have the following result, which is proved in S1 Text §2.5.

##### Theorem 3.

*Given a network* 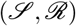. *Let* 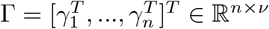 *be its stoichiometry matrix. If the following algorithm terminates successfully, then* 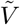 *is an RLF*.

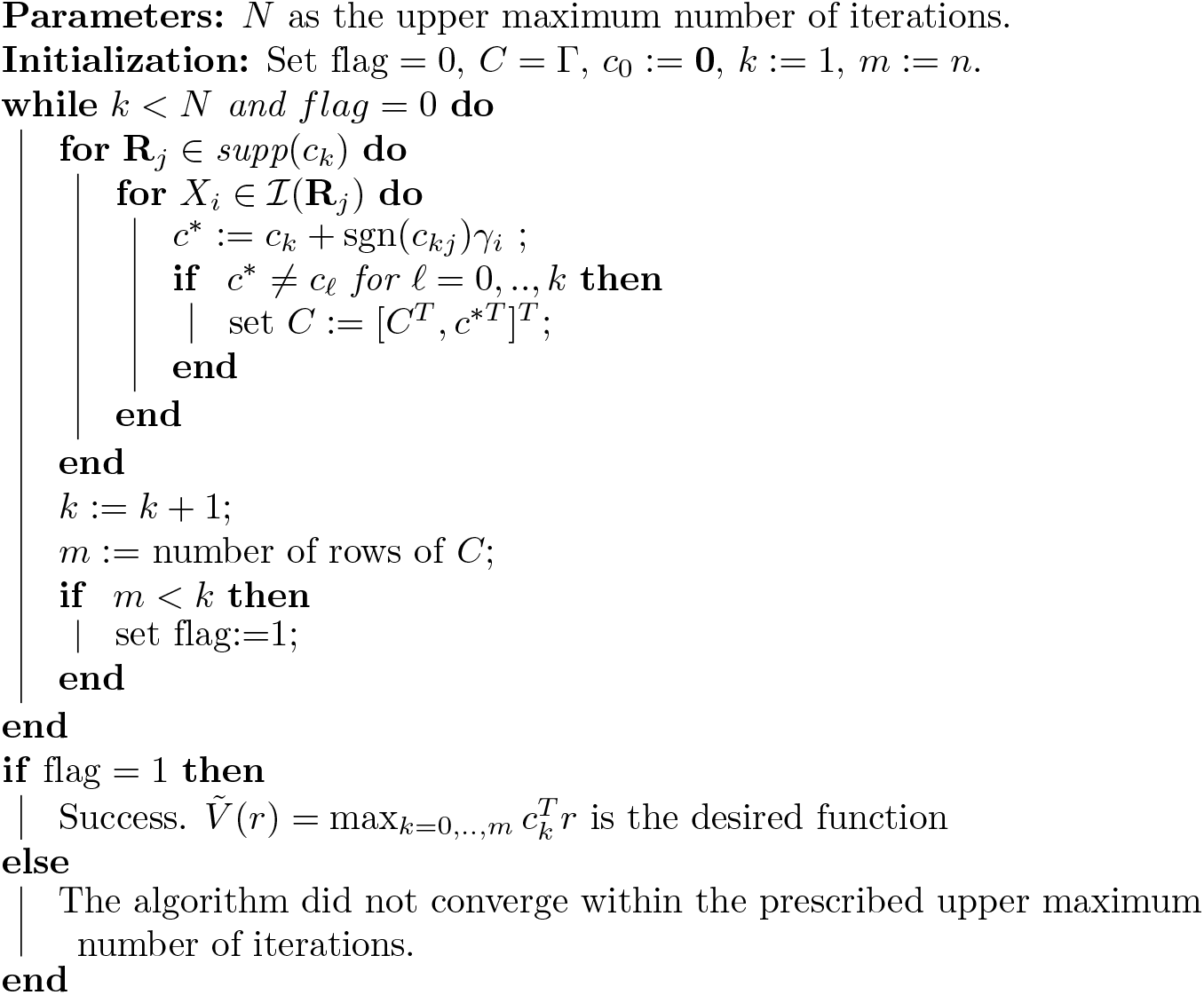

The algorithm above is computationally very light compared to the linear program with a large *H*. Furthermore, if the network satisfies AS1 then the RLF can be written as 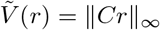.

#### Graphical criteria for the construction of Max-Min RLFs

Compared to computational conditions, it is highly desirable to have graphical conditions and some have been provided in [26, 27]. We reformulate those conditions to be more friendly for computational implementation in `LEARN`. Those conditions enable the identification of attractive networks by mere inspection of the reaction graph for a particular class of networks.

We introduce some notations. Let 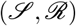 be a given *non-autocatalytic* network that satisfies AS1. Consider the decomposition 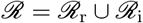 into the subsets of reactions that are reversible and irreversible, respectively. Furthermore, we can decompose 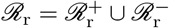 into the forward and backward reactions, respectively. Let 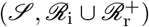 be the corresponding *irreversible subnetwork* and let 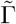 be its stoichiometry matrix. Since the designation of a forward and reverse reaction is arbitrary, we need a decomposition such that 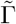 has a one-dimensional nullspace. If such a decomposition exists, then we call the original network 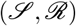 an *M-network*. Our graphical condition applies to this class of networks, and it can be stated as follows.

##### Theorem 4.

*Let* 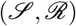 *be an M-network, and let* 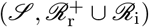 *be the subnetwork defined above, where the reactions are enumerated as* 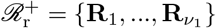, 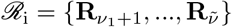. *If the irreversible subnetwork satisfies the following properties:*

1. *each species participates in exactly one reaction, and*
2. *each reaction* 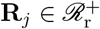 *satisfies the following statement: If a species X*_*i*_ *is a product of* **R**_*j*_, *then X*_*i*_ *is not a product of another reaction, then*

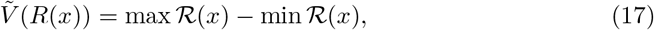

*where* 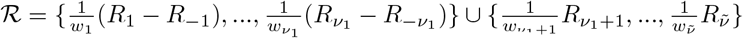, *is a convex PWL RLF, where* 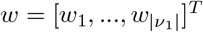 *belongs to the null space of* 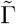.

#### Piecewise Quadratic-in-Rates RLFs

The framework developed in this paper allows us to go beyond PWL RLFs, and consider other classes of functions such piecewise quadratic-in-rate functions of the form:

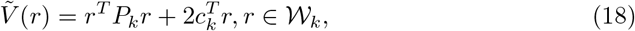

for some matrices *P_k_* ∈ ℝ^*ν*×*ν*^, *c_k_* ∈ ℝ^*ν*^, *k* = 1, ‥, *m*.

Instead of linear programming, construction of PWQR RLFs is a copositive programming problem. Although copositive programs are convex, solving them generally is shown to be NP-hard [94]. Therefore, we use a common relaxation scheme based on the observation that the class of copositive matrices encompasses the classes of positive semi-definite matrices, and nonnegative matrices. The following theorem states the result and it is proven in S1 Text §3.1.

##### Theorem 5.

*Given a network* 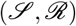 *that satisfies AS1 and a partitioning matrix H* ∈ ℝ^*p*×*r*^. *Let* {*v_i_*} *be the basis for the kernel of* Γ *Consider the following semi-definite program:*

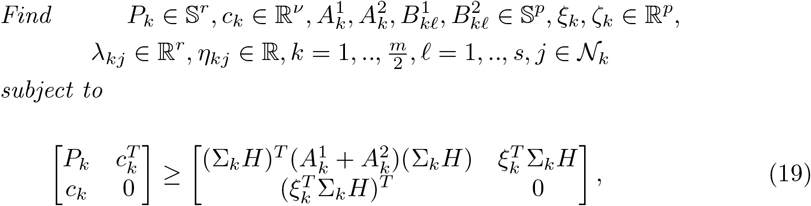

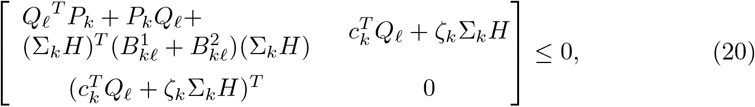

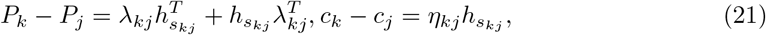

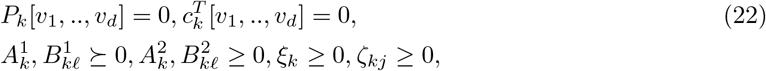

*where d* = dim(ker Γ) *and* 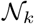 *is the set of neighbor of region* 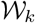 *(see S1 Text §3.2). If the SDP is feasible, then* 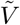 *as defined in* (18) *is an RLF for* 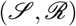 *if* 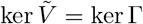.

This class of networks for which PWQ RLFs exist is potentially larger than that of PWL RLFs even when we set *c_k_* = 0, *k* = 1, ‥, *m* in (18) as the following proposition establishes. The proof is given in S1 Text §X.

##### Proposition 6.

*Let a network* 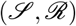 *that satisfies AS1 be given. If there exists an RLF* 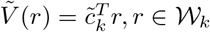 *with a partition matrix H, then the SDP problem in Theorem 5 with* 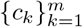 *constrained to be zeros is feasible. In particular*, 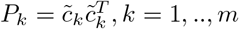 *is a feasible solution*.

### Properties of Attractive Networks

#### Robust Non-degeneracy

It has been shown in [27] that the negative Jacobian of any network admitting a PWL RLF is *P*_0_, which means that all principal minors are nonnegative. We show that the reduced Jacobian (i.e, Jacobian with respect to a stoichiometric class) is non-degenerate for *all admissible kinetics* if it is so at one interior point only. The proof is stated in S1 Text §4.1.

##### Theorem 7.

*Assume that there exists a PWL RLF. If for some kinetics* 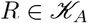 *there exists a point in the interior of a proper stoichiometric class such that the reduced Jacobian is non-singular at it, then the reduced Jacobian is non-singular in the interior of* 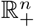 *for all admissible kinetics*.

In `LEARN`, robust non-degeneracy is checked with *ρ_ℓ_* = 1, *ℓ* = 1, …, *s*. It amounts to checking the non-singularity of one matrix.

##### Remark 2.

*Robust non-degeneracy, coupled with the existence of a PWL RLF, automatically guarantees the uniqueness of positive steady states and their exponential stability (see S1 Text §4.1.2,§4.1.3). Globally stability has been checked via a LaSalle algorithm in [27], which is automatically satisfied for conservative M-networks. Alternatively, global stability follows automatically for any positive steady state if the network is robustly nondegenerate [95]. Hence, Theorem 7 can be used to verify global stability when a PWL RLF exists. Note, however, that the test above is with respect to the stoichiometric class only. In the case of degenerate reduced Jacobians, a stoichiometric class can be partitioned further into* kinetic compatibility classes *[16]. The graphical LaSalle’s algorithm applies to such cases also*.

#### Absence of Critical Siphons

A siphon is any (minimal) set of species which has the following property: if those species start at zero concentration, then they stay so during the course of the reaction [41]. Siphons are of two types: trivial and critical. A trivial siphon is a siphon that contains the support of a conservation law. A critical siphon is a siphon which is not trivial. Critical siphons can be found easily from the network graph. The absence of critical siphons in a network has been shown to imply that it is *structurally persistent* (for conservative networks or systems with bounded flows) [41]. Informally, a system is persistent if the following holds: if all species are initialized at nonzero concentrations, none of them will become asymptotically extinct. We show that the existence of critical siphons precludes the existence of RLF under mild conditions which serves as an easy-to-check condition to preclude the existence of an RLF. Review of the concept of siphons and the proof the result is included in S1 Text §4.2.

##### Theorem 8.

*Given a network* 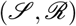 *that satisfies AS1. Assume it has a critical siphon* 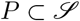. *Let* 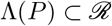 *be the set of reactions for which the species in P are reactants. Then there cannot exist a PWL RLF if any of the following holds:*

1. 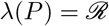, *i.e P is a critical deadlock*.
2. 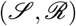 *is a conservative M network*.
3. 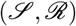 *is conservative and has a positive non-degenerate steady state for some admissible kinetics*.

##### Remark 3.

*The tests established in Theorem 8 have been implemented in `LEARN`*.

### RLFs in Other Coordinates

In this subsection we study an alternative RLF and we link the results with the ones proposed in [34, 36]. We will show that any RLF has an alternative form if it satisfies a mild condition. In particular, all PWL RLFs have alternative forms. Assume that (1) has a steady state *x_e_*. Then, we ask whether there exists a Lyapunov function of the form 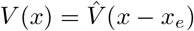. However, note that this Lyapunov function decreases only in the stoichiometric class containing *x_e_* and that computing its level sets requires knowing *x_e_*. We call 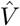 a *concentration-dependent RLF*. Similar to before, we will characterize the existence of an RLF of the form 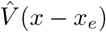 for a network 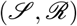 by the existence of a common Lyapunov function for a set of extremals of an appropriate cone. In this subsection, we assume that there exists a positive 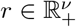 such that Γ*r* = 0.

We will adopt an alternative representation of the system dynamics. Consider a CRN as in (1), and let *x_e_* be a steady state. Then, there exists 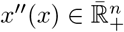 such that (1) can written equivalently as:

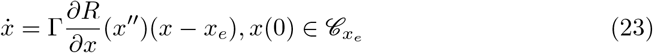

The existence of *x″* ≔ *x_e_* + *ε_x_*(*x* − *x_e_*) for some *ε_x_* ∈ [0, 1] follows by applying the Mean-Value Theorem to *R*(*x*) along the segment joining *x_e_* and *x*.

Similar to the analysis for a rate-dependent RLFs, the Jacobian of (1) can be shown to belong to the conic span of a set of rank-one matrices 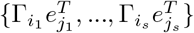 where {Γ_1_, ‥, Γ_*n*_} are the columns of Γ. The pairs (*i_ℓ_*, *j_ℓ_*), *ℓ* = 1, ‥, *s* are the same pairs used before.

Let *D^T^* be a matrix with columns that are the basis vectors of ker Γ^*T*^. The following theorem is proven in S1 Text §5.1.

#### Theorem 9.

*Given a network* 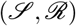. *There exists a common Lyapunov function* 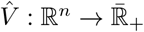 *for the set of linear systems* 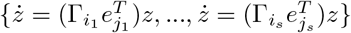, *on the invariant subspace* 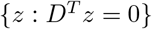 *if and only if* 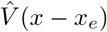 *is a concentration-dependent RLF for any x*_*e*_.

#### Relationship between the RLFs in concentration and rates

We show next that if 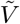 is a rate-dependent RLF that satisfies a relatively mild additional assumption, then then 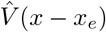 is a concentration-dependent RLF, where *x_e_* is a steady state point for (1). The following theorem can be stated and is proved in S1 Text §5.2.

##### Theorem 10.

*Let* 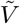 *be an RLF for the network* 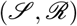. *If there exists* 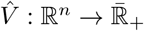 *such that for all r* ∈ ℝ^*ν*^:

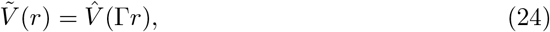

*then* 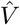 *is a concentration-dependent RLF for the same network*.

#### PWL functions in concentrations

All PWL RLFs constructed before have the property that there exists 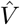 such that 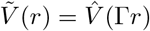. Hence, there exists a concentration-dependent PWL RLF for the same network. In particular, consider a PWL RLF defined with a partitioning matrix *H* as in (16). By AS1 and the assumption that ker *H* = ker Γ, there exists *G* ∈ ℝ^*p*×*n*^ and 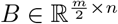 such that *H* = *G*Γ and *C* = *B*Γ. Similar to 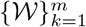, we can define the regions:

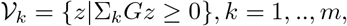

where it can be seen that 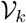 has nonempty interior iff 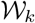 has nonempty interior.

Therefore, as the pair (*C, H*) specifies a PWL RLF, the pair (*B, G*) also specifies the function:

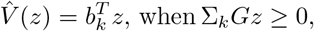

where 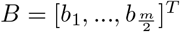. If 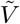 is convex, then it can be written in the form: *V*_1_(*x*) = ‖*CR*(*x*)‖_∞_. Similarly, the convexity of 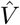 implies that *V*_2_(*x*) = ‖*B*(*x* − *x_e_*)‖_∞_, where the latter is the Lyapunov function used in [36].

Theorem 10 shows how to go from a rate-dependent to a concentration-dependent RLF. The following theorem shows that one can start with either PWL RLF to get the other. It is proved in S1 Text §5.3.

##### Theorem 11.

*Given* 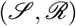. *Then, if*

1. (*B*Γ, *G*Γ) *specifies a rate-dependent PWL RLF, then* (*B, G*) *specifies a concentration-dependent PWL RLF*.
2. (*B, G*) *specifies a concentration-dependent PWL RLF, then* (*B*Γ, *G*Γ) *specifies a rate-dependent PWL RLF*.

##### Remark 4.

*Since D^T^* (*x* − *x_e_*) = 0 *for* 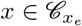, *then if* ‖*B*(*x* − *x_e_*)‖_∞_ *is an RLF, then* ‖(*B* + *Y D^T^*)(*x* − *x_e_*)‖_∞_ *is also an RLF for an arbitrary matrix Y. Furthermore, since Theorem 11 has shown that the concentration-based and the rate-based representations are equivalent, it is easier to check and construct RLFs in the rate-based formulation and they hold the advantage of being decreasing for all trajectories over all stoichiometry classes*.

#### Computational Package

Calculations were performed using MATLAB 10 via our software package `LEARN` available at https://github.com/malirdwi/LEARN. Available subroutines and example runs are included in S1 Text §7. The package `cvx` [96] has been used for solving linear and semi-definite programs, and the package PetriBaR for enumerating siphons [97].

## Supporting Information Legend

**S1 Text** Supporting information file with mathematical proofs, generalization of the results and additional information.

## Supplementary Information

### 1 Lyapunov’s Second Method

#### 1.1 Preliminaries

First, let consider *specific* kinetics 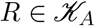. Hence, the ODE is given as

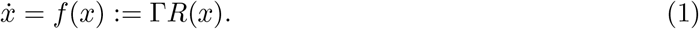

We have the following definition:

##### Definition A-1.

*Given the ODE* (1). *Let* 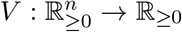 *be locally Lipschitz. Then V is said to be a Lyapunov function for* (1) *if V is*

- Positive-Definite *(with respect to the steady states set) if V* (*x*) ≥ 0, *and V* (*x*) = 0 *if and only if R*(*x*) ∈ ker Γ.
- Nonincreasing *if* 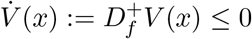 *for all x, where* 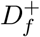 *is defined below*.

Note that when *∂V/∂x* exists at a point *x*, then 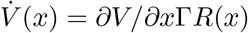.

##### 1.1.1 Generalized Derivatives

The function *V* is locally Lipschitz, and hence it is not necessarily differentiable everywhere. It has been known since the early stability literature (see [1], [2]) that the standard Lyapunov theorems can be generalized without difficulty with locally Lipschitz Lyapunov functions and Dini derivatives.

The upper Dini derivative for *V* in the direction of a function *f* (*x*) ≔ Γ*R*(*x*) is defined as:

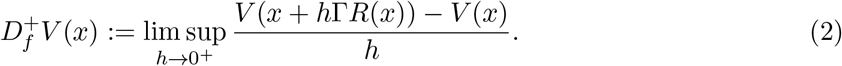

For a locally Lipschitz function, the above quantity is always finite.

An alternative definition of the derivative, which is more restrictive but has more convenient calculus, is the *Clarke derivative*, which is defined as [3]:

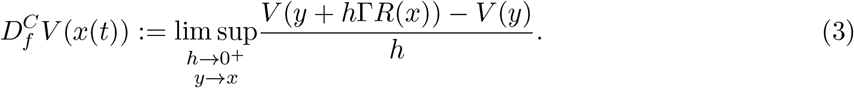

Note that 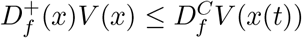. We will define 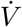 in the sense of Dini.

##### 1.1.2 LaSalle’s Condition

Conventional stability theory [1] examines stability with respect to an isolated steady state. However, for reaction networks, there is usually a *continuum* of equilibria. This means that asymptotic stability or Lyapunov stability are not achieved in the classical sense. Nevertheless, the state space of reaction networks is divided into stoichiometric compatibility classes which are forward invariant. Furthermore, a stoichiometric class can sometimes be divided into *kinetic compatibility classes* [4]. In general, any initial condition *x*_◦_ is associated with a compatibility class 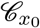. Hence we state the following definition:

###### Definition A-2 (The LaSalle’s Condition).

*Given an ODE* (1) *with a Lyapunov function V. The* LaSalle’s Condition *is satisfied if the following statement holds:*

*If a solution φ*(*t*; *x*_◦_) *of* (1) *satisfies* 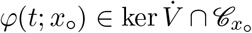 *for all t* ≥ 0, *then* 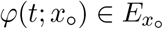 *for all t* ≥ 0, *where* 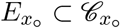 *is the set of steady states for* (1) *contained in* 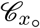.

##### 1.1.3 Lyapunov Stability Theorem

We state the following theorem which is standard Lyapunov theory adapted to our settings [5].

###### Theorem A-1 (Lyapunov’s Second Method).

*Given* (1) *with initial condition* 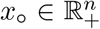. *Let* 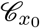 *be its class. Assume there exists a Lyapunov function V and suppose that x*(*t*) *is bounded*.

- *Then the steady state set* 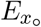 *is Lyapunov stable relative to* 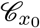.
- *If, in addition, the LaSalle’s Condition is satisfied, then* 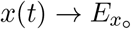 *as t* → ∞ *(i.e., the point to set distance of x*(*t*) *to* 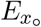 *tends to* 0*). Furthermore, any isolated steady state relative to* 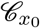 *is asymptotically stable*.
- *If the LaSalle’s condition is satisfied, and all the trajectories are bounded, then: if there exists an* 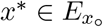 *which is isolated relative to* 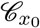, *then it is unique, i.e.*, 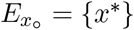. *Furthermore, it is globally asymptotically stable steady state relative to* 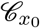.

#### 1.2 Robust Lyapunov Functions and Proof of Theorem 1

In the main text, we have defined an RLF 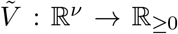. For a given 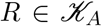, the Lyapunov function is 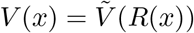.

Before proving Theorem 1, we need to state and prove the following Lemma:

##### Lemma A-1.

*Let* 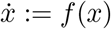, *and let* 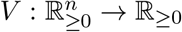 *be a locally Lipschitz function such that:*

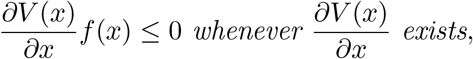

*Then* 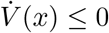 *for all x*.

*Proof*. Since *V* is assumed to be locally Lipschitz, Rademacher’s Theorem implies that it is differentiable (i.e., gradient exists) almost everywhere [3]. Recall that for a locally Lipschitz function the *Clarke gradient* at *x* is defined as *∂_C_ V* (*x*) ≔ co *∂V* (*x*), where:

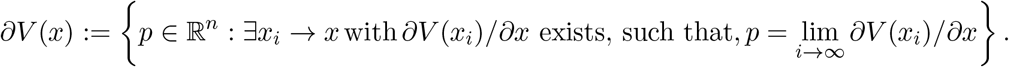

Let *p* ∈ *∂V* (*x*) and let 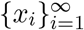 be any sequence as in the definition of the Clarke gradient. By the assumption stated in the Lemma, (*∂V*(*x_i_*)/*∂x*)*f*(*x_i_*) ≤ 0, for all *i*. Hence, the definition of *p* implies that *p*^*T*^ *f* (*x*) ≤ 0. Since *p* was arbitrary, the inequality holds for all *p* ∈ *∂V* (*x*).

Now, let 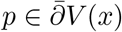 where *p* = ∑_*i*_ *λ*_*i*_*p*_*i*_ is a convex combination of any *p*_1_, …, *p*_*n*+1_ ∈ *∂V* (*x*). By the inequality above, 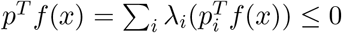. Hence *p*^*T*^*f*(*x*) ≤ 0 for all 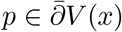. As in [3], the Clarke derivative of *V* at *x* in the direction of *f*(*x*) can be written as 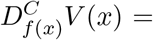 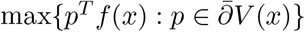. By the above inequality, we get 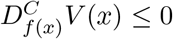 for all *x*. Since the Dini derivative is upper bounded by the Clarke derivative, we finally get:

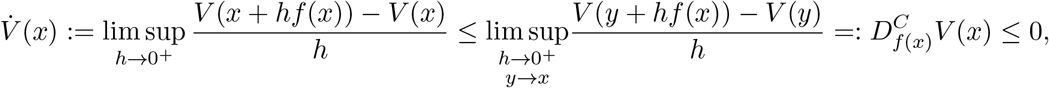

for all *x*.

##### Proof of Theorem 1

We show that the existence of the common Lyapunov function implies the existence of the RLF. Nonnegativity of *V* follows from the nonnegativity of 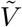. Let (*i, j*) = *κ*(*ℓ*), and recall that 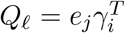, hence 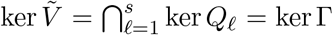. Therefore, *R*(*x*) ∈ ker *V* iff Γ*R*(*x*) = 0, which establishes the positive-definiteness of *V*.

We assumed that 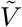 has a negative semi-definite time-derivative for every linear system in the considered set. Hence, whenever 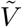 is differentiable at a point *r*, we can write 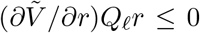, 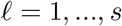. Hence, for any 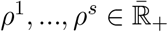:

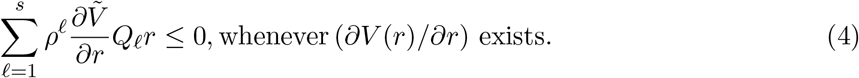

Therefore, whenever 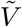 is differentiable we have

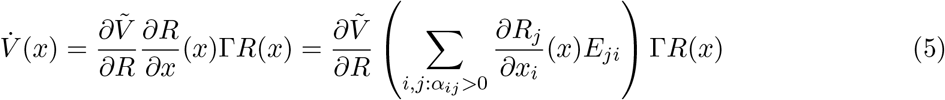

where 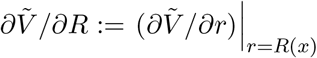.

Now, denote 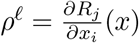, which is nonnegative by AK3. This allows us to write:

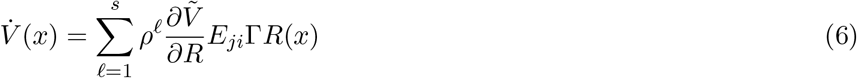

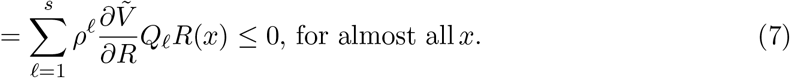

The last inequality follows from (4). Using Lemma A-1, 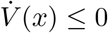 for all *x*, and for all 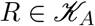.

In order to show the other direction, since most of the properties outlined in the RLF definition are clearly satisfied, it remains to show nonincreasingness. Assume that there exists *ℓ* such that 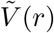 is not nonincreasing along the trajectories of 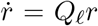. Consider the corresponding term in (6). Since *V* (*R*(*x*)) is a Lyapunov function for any choice of admissible rate reaction function *R*, choose 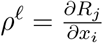 to be large enough such that 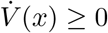 for some *x*; this results in a contradiction.

### 2 Piecewise Linear RLFs

#### 2.1 Checking a candidate Lyapunov function

Suppose we are given a matrix *H* ∈ ℝ^*p*×*r*^ such that ker *H* = ker Γ. Let 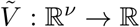 be a continuous PWL function given as

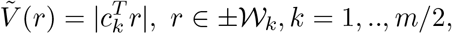

where the regions 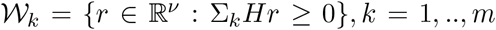 form a proper conic partition of ℝ^*ν*^, while 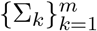 are signature matrices (diagonal matrices with ±1 on the diagonal) with the property Σ_*k*_ = −Σ_*m*+1−*k*_, *k* = 1, ‥, *m/*2. (see [5] for a detailed exposition on the geometry of the partition regions).

The coefficient vectors of each linear component can be collected in a matrix 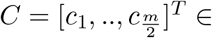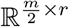. If the function 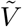 is convex, then we have the following simplified representation of *V* [6]:

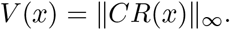

This representation is analogous of the *ℓ*_∞_-norm Lyapunov functions that have been used for linear systems in [7].

##### Theorem A-2.

*Let* Γ *and H be given as above. Let* 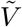 *be a candidate continuous nonnegative PWL function with* 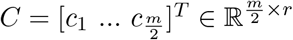. *Then* 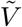 *is an RLF if and only if:*

- ker *C* = ker Γ, *and*
- *there exists matrices* 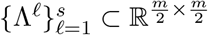 *such that*

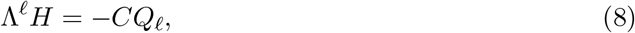

*and* 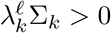, *where* 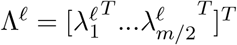. *If* 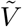 *is convex, then the second condition can be replaced with* *2) there exists Metzler matrices* 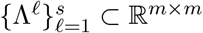 *such that*

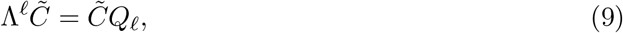

*and* Λ^*ℓ*^**1** = 0 *for all ℓ* = 1, ‥, *s, where* 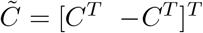.

*Proof*. The proof can be carried out by performing algebraic manipulations on the results presented in [5, Theorem 4]. The function has been assumed continuous and nonnegative. It remains to show that the corresponding condition in [5] is equivalent to (9). Considering (9) row by row, it can be written in the following form for 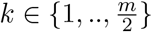:

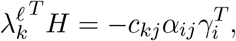

where (*i, j*) = *κ*(*ℓ*). *c_kj_* can be replaced with sgn(*c_kj_*) and *α_ij_* can be replaced with 1 if we are considering only *i* ∈ *I_k_*, *j* ∈ *J_ki_*. Therefore, equivalence with the corresponding condition in [5, Theorem 4] is established.

For a convex PWL function, we can write the corresponding condition in [5, Theorem 5] as follows after replacing sgn(*c_kj_*) by *c_kj_*, and inserting *α_ij_*:

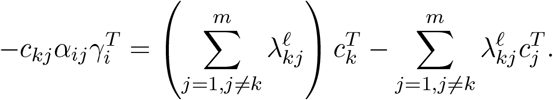

Let 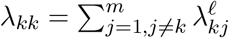. Then

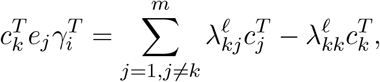

which enforces Λ^*ℓ*^ to be Metzler and Λ^*ℓ*^1 = 0 as above.

##### Remark A-1.

*The symmetries in equation* (9) *imply that it can be written equivalently as:*

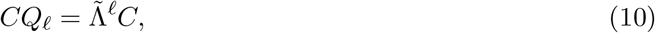

*where* 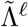 *is an* 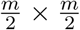 *matrix which is defined by subtracting the upper* 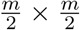 *blocks of* Λ^*ℓ*^ *from each other. The matrix* 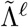 *satisfies:*

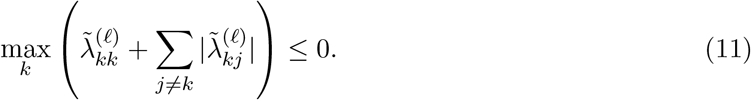

*This is exactly the condition that ℓ*_∞_*-norm Lyapunov functions need to satisfy for a linear system [8, 9]. This shows that Theorem 1 provides the framework to utilize the existing linear stability analysis techniques in the literature to construct robust Lyapunov functions for nonlinear systems such as CRNs. For example, we can verify ℓ*_1_ *Lyapunov functions of the form V* (*x*) = ‖*CR*(*x*)‖_1_ *directly by replacing condition* (11) *by*

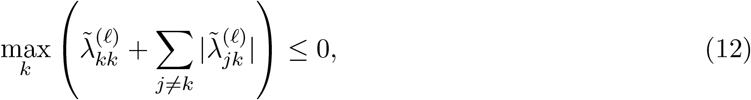

*instead of converting them to the ℓ*_∞_*-norm form*.

#### 2.2 Proof of Theorem 2

The linear program has the parametrization 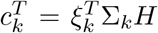, which follows from applying Farkas’s Lemma to ensure that 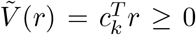 on the region 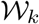. (See [5] for full details.) The second condition: *CQ*_*ℓ*_ = −*Q*_*ℓ*_*H* follows from Theorem 2 and ensures that *V* (*R*(*x*)) is nonincreasing. To ensure continuity we need to have 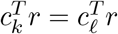 whenever 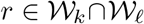. Since ker *H* = ker Γ, continuity can be imposed by the constraint (*c_k_* − *c*_*ℓ*_)^*T*^ *v_i_* = 0 for *i* = 1, ‥, dim(ker Γ).

#### 2.3 Enforcing convexity in a linear program

The linear program presented in the main text does not enforce convexity on the PWL RLF. Following [5], we describe how to write a linear program to construct convex PWL RLFs in what follows.

We need to introduce the concept of a neighbor to a region. Fix *k* ∈ {1, ‥, *m/*2}. Consider the matrix *H*: for any pair of linearly dependent rows 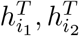 eliminate 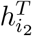. Denote the resulting matrix by 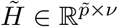, and let 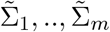 the corresponding signature matrices. Therefore, the region can be represented as 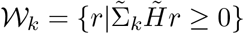. The *distance d_r_* between two regions 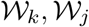 is defined to be the *Hamming distance* between 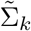 and 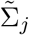. Hence, the set of neighbors of a region 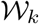 are defined as:

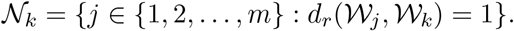

Equivalently, note that a neighboring region to 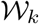 is one which differs only by the switching of one inequality. Denote the index of the switched inequality by the map 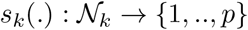. For simplicity, we use the notation *s_kℓ_* ≔ *s_k_*(*ℓ*).

##### Theorem A-3.

*Given the system* (1) *and a partitioning matrix H* ∈ ℝ^*p*×*r*^. *Consider the linear program:*

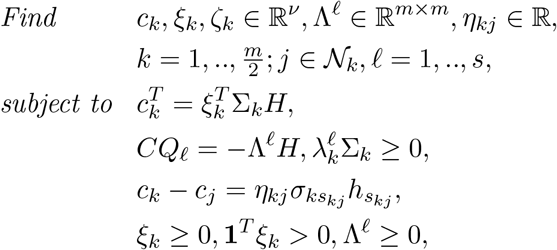

*where σ_kj_ is the jth entry on the diagonal of* Σ_*k*_. *Then there exists a PWL RLF with partitioning matrix H if and only if there exists a feasible solution to the above linear program that satisfies* ker *C* = ker Γ. *Furthermore, the PWL RLF can be made convex by adding the constraints η*_*kj*_ ≥ 0.

#### 2.4 Networks without positive steady states

Let AS1 be the assumption that requires the existence of a positive vector in ker Γ, which is a necessary condition for the existence of positive steady states. This assumption simplifies the geometry of the partition regions and enforces symmetry on the coefficient matrix *C*. Nevertheless, our techniques can be extended without difficulty for the construction of PWL RLFs for generic CRNs that do not satisfy AS1. Consider a matrix *H* ∈ ℝ^*p*×*ν*^, with ker *H* = ker Γ. The regions are defined as:

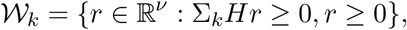

where *k* = 1, ‥, 2^*p*^. Note that the inequality *r* ≥ 0 needs to be explicitly included. As before, let *m* be the number of non-empty interior regions. Then, the regions are ordered such that the first *m* regions are the non-empty interior ones. Therefore, the following theorem can be stated for networks that do not necessarily satisfy AS1.

##### Theorem A-4.

*Consider the system* (1), *with* 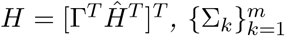 *given as before. Consider the following linear program:*

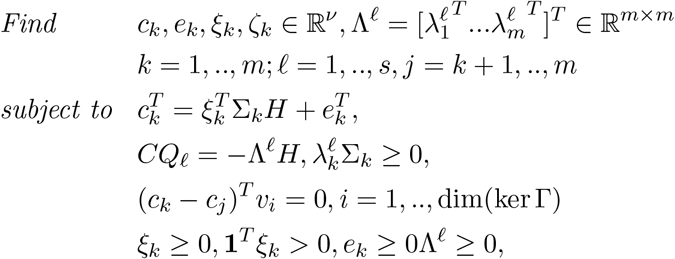

*Then there exists a PWL RLF with partitioning matrix H if and only if there exists a feasible solution to the above linear program with* ker *C* = ker Γ *satisfied*.

#### 2.5 Proof of Theorem 3

The algorithm starts with *C* = Γ. Hence, it can be interpreted as an initial PWL function 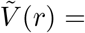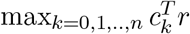 where *c_k_* = *γ_k_*, *k* = 1, ‥, *n*.

We aim at restricting the *active* region of each function 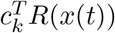 to the region on which it is nonincreasing, i.e 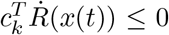. This is accomplished by adding extra linear components that ensures this. Define the *active region* of a vector *c_k_*, *k* = 1, ‥, *m*_0_, as:

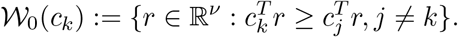

We define the *permissible region* of a linear component *c_k_* to be the region for which it is nonincreasing:

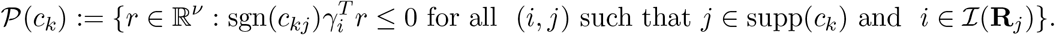

Note that in general, 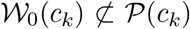. Therefore, the iterative procedure defines a new PWL function with matrix *C*_1_ so that 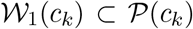. To achieve this, new rows are added to *C* as follows:

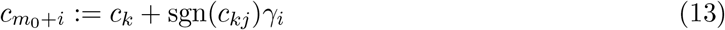

for all (*i, j*) such that *j* ∈ supp(*c_k_*) and 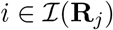.

The procedure is repeated for every row of *C*. If the procedure terminates, i.e no new rows need to be added, then 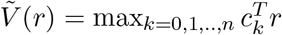 is a PWL RLF.

### 3 Construction of PWQ RLFs

#### 3.1 Proof of Theorem 5

We show that 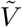 is a common Lyapunov function for 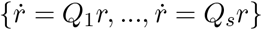 as in Theorem 1. In order to show nonnegativity, inequality (23) implies that:

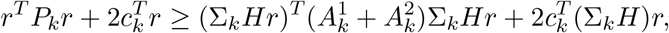

and since 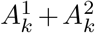 is copositive this implies that 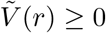 when 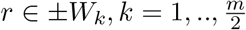, which establishes nonnegativity. Positive-definiteness follows from (26) and the assumption in the statement of the theorem.

For continuity, it is sufficient to establish it between neighboring regions. Therefore, assume 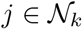, and let 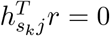 be the intersection hypersurface, then (25) implies that 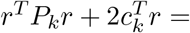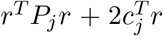 when 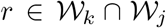. The other direction holds also by writing *P_k_* − *P_j_* over the decomposition 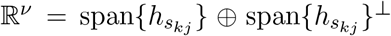. Continuity of 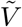 implies also that 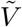 is locally Lipschitz.

In order to show that this derivative is negative semi-definite, consider the **ℓ**^th^ system, and let 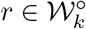, then:

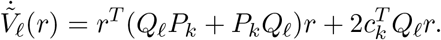

Note that (24) implies that 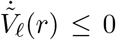 when 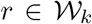. As it is true for all *k*, then 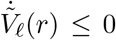 for all *ℓ* = 1, ‥*s*, and all r such that 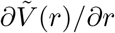 exists. By Lemma 1, this implies that 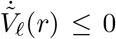 for *ℓ* = 1, ‥, *s*.

#### 3.2 Proof of Proposition 6

Assume that 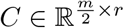 is given such that 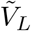 is a PWL RLF, where 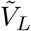 is defined as in (2.1). Let 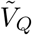 be defined by

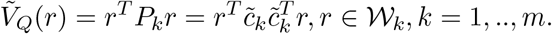

The constraints (25),(26) are clearly satisfied. The inequality (23) is satisfied with 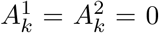, 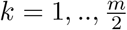. It remains to show that (24) is satisfied.

Fix *ℓ* ∈ {1, ‥, *s*}, 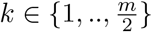. Then

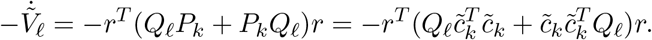

Since it is assumed that 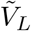 is PWL RLF, there exist 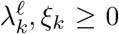 such that 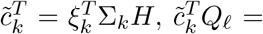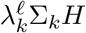. Therefore:

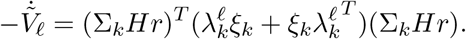

Hence, (24) is satisfied with 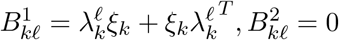.

### 4 Properties of Attractive Networks

#### 4.1 Robust non-degeneracy

A point *x_e_* of (1) is *non-degenerate* if the Jacobian evaluated at *x_e_* relative to 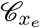 is nonsingular. More precisely, let us change coordinates using a transformation matrix 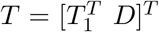, where *D^T^* has full row rank and *D^T^*Γ = 0, and *T*_1_ is any matrix such that *T* is nonsingular. Then, the Jacobian in the new coordinates can be written as:

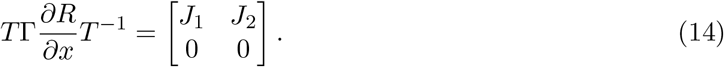

Therefore, *x_e_* is nondegenerate iff *J*_1_ evaluated at *x_e_* is nonsingular. The matrix *J*_1_ is called a *reduced Jacobian*.

##### 4.1.1 Proof of Theorem 7

Recall that for an attractive network with PWL RLF, the negative Jacobian is *P*_0_ for any choice of 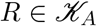 [5]. Using the Cauchy-Binet formula [10], let *I* ⊂ {1, ‥, *n*} be an arbitrary subset so that |*I*| = *k*. The corresponding principal minor can be written as:

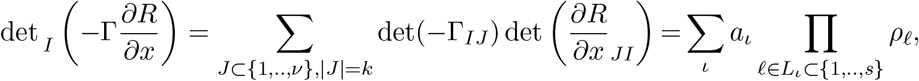

where the last equality refers to the fact that the sum can be expressed as a linear combination of products of *ρ*_1_, …, *ρ_s_*. We claim that the coefficients *a_ι_* are all nonnegative. To show this, assume for the sake of contradiction that there is some negative 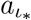. If we set all *ρ*’s to zero except the ones appearing in the 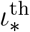 term, then this implies that the corresponding principal minor can be negative; a contradiction.

Now, the theorem can be proven by noting that the reduced Jacobian is non-singular iff the sum of all *k* × *k* principal minors of the negative Jacobian is positive, where *k* = rank(Γ). Since it is assumed that there exists a point for which the reduced Jacobian is non-singular, this implies that the sum of principal minors is positive for some choice of *ρ*_1_, ‥, *ρ_s_*. Since all of the principal minors are nonnegative, then at least one of them is positive. By AK4, that principal minor stays positive for any choice of positive *ρ*_1_, ‥, *ρ_s_*, i.e. it stays positive over the interior of 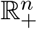.

##### 4.1.2 Uniqueness of the steady states

The following directly from Theorem 7.

###### Proposition A-5.

*Consider a network* 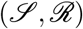 *that satisfies AS1 and admits a PWL RLF. If there exists a non-degenerate positive steady state x*_*e*_, *relative to* 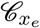, then it is unique.

*Proof*. Theorem 7 has shown that the existence of an non-degenerate positive steady state *x_e_* ensures that the reduced Jacobian is non-singular on the interior of the orthant. In order to show uniqueness, assume for the sake of contradiction that there exists *y* ≠ *x_e_*, 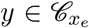 such that Γ*R*(*y*) = 0. Then the fundamental theorem of calculus implies,

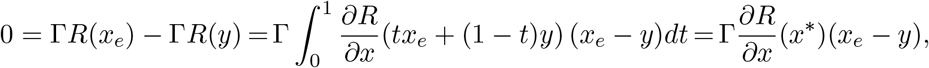

where *x*^∗^ = *t*^∗^*x_e_* +(1−*t*^∗^)*y*, and *t*^∗^ ∈ (0, 1). The existence of *t*^∗^ is implied by the integral mean-value theorem. Since 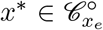, then the reduced Jacobian at *x*^∗^ is non-singular relative to Im Γ. Since *x_e_* − *y* ∈ Im Γ, then *y* = *x_e_*. This gives a contradiction.

##### 4.1.3 Exponential stability

We have shown that the existence of a PWL RLF function implies that it is a common Lyapunov function for all linear systems that belong to a linear differential inclusion.

In fact, one of the properties of systems that admits a piecewise linear Lyapunov function is that a stable steady state cannot have purely imaginary eigenvalues [11]. Hence, the reduced Jacobian at a non-degenerate steady state cannot admit pure imaginary eigenvalues which implies the following Theorem:

###### Theorem A-6.

*Let* 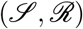 *be a network that admits a PWL RLF. If a positive steady state x*_*e*_ *is non-degenerate relative to* 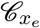, *then it is exponentially asymptotically stable*.

##### 4.1.4 Global stability

Establishing global asymptotic stability of a positive steady state for a network that admits a PWL RLF has been accomplished via a LaSalle graphical algorithm in [5]. Nevertheless, if a network is known to be robustly non-degenerate with respect to the stoichiometric class (by the test given in Theorem 7 for instance), then the following result holds:

###### Theorem A-7.

*([12]) Suppose that the system* (1) *admits a PWL RLF. If the Jacobian is robustly non-degenerate relative to a stoichiometric class* 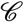, *then every positive steady state* 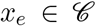 *is globally asymptotically stable relative to* 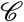.

Hence, the graphical LaSalle algorithm is not needed for networks with Jacobians that are robustly non-degenerate relative to stoichiometric classes.

#### 4.2 Absence of critical siphons: Proof of Theorem 8

Assume *P* is a critical siphon for the Petri-net associated with Γ, and let *n_p_* = |*P*|. Let Λ(*P*) be the set of output reactions of *P*, and let *ν_p_* = |Λ(*P*)|.

Item 1 of Theorem 8 has been proved in [5]. We restate the proof in this paper’s terminology for completeness. First, the following lemma is needed.

##### Lemma A-2.

*Consider a network* 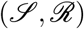. *Let P be a set of species that does not contain the support of a conservation law; let its indices be numbered as* {1, …, *n_p_*}. *Then, there exists a nonempty-interior region* {*r*|Σ_*k*_Γ*r* ≥ 0} *with a signature matrix* Σ_*k*_ that satisfies 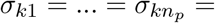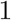.

*Proof*. Assume the contrary. This implies that 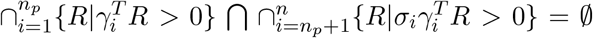 for all possible choices of signs *σ*_*i*_ = ±1. However, ℝ^*r*^ can be partitioned into a union of all possible half-spaces of the form 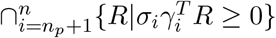. Therefore, this implies that 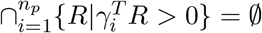. By Farkas Lemma, this implies that there exists 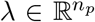 satisfying *λ* > 0 such that [*λ^T^* 0]Γ = 0. Therefore, *P* contains the support of the conservation law [*λ^T^* 0]^*T*^ ; a contradiction.

Therefore, we can state the proof of the first item:

##### Proof of Theorem 8-1).

Without loss of generality, let {1, …, *n_p_*} be the indices of the species in *P*. Using Lemma 2, there exists a nonempty-interior sign region 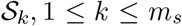 with a signature matrix Σ_*k*_ that satisfies 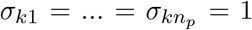. Since 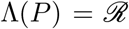, this implies that we must have *c_k_* ≥ 0 to match the sign pattern of Σ_*k*_. But since ∃*v* ≫ 0 ∈ ker Γ, then this implies that *c_k_* = 0 which contradicts the positive definiteness condition on the RLF since ker *C* ≠ ker Γ.

In order to proceed, we denote by Ψ_*P*_ the *face* that corresponds to a siphon *P*. It is given by: 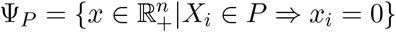. We state the following lemma next:

##### Lemma A-3.

*Consider a network* 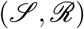. *Let P be a critical siphon, and let* Ψ_*P*_ *be the associated face. If the network is conservative, then for any proper stoichiometric compatibility* 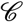, *there exists a steady state x*_*e*_ *of* (1) *such that* 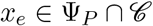.

*Proof*. The set 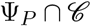 is compact, forward invariant, and convex, since both sets Ψ_*P*_, 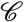 are such. Hence, the statement of the lemma follows directly from the application of the Brouwer Fixed Point Theorem on the associated flow.

We are ready now to prove the second item of Theorem 8.

##### Proof of Theorem 8-2).

By Lemma 3, there exists a steady state in Ψ_*P*_. Since it is assumed that there exists a non-degenerate steady state in the interior, Proposition 5 implies that the network cannot admit a PWL RLF.

Before concluding the proof, a simple lemma is stated and proved:

##### Lemma A-4.

*Let x*_*e*_ *be a steady state of* (1). *Let* 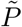 *be a set of species that correspond to* {1, ‥, *n*}\ *supp*(*x_e_*). *Then*, 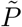 *is a siphon*.

*Proof*. Assume that 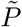 is not a siphon, then there exists some 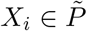 and 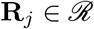 such that *X_i_* is a product of **R**_*j*_ and 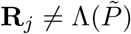. At the given steady state, all negative terms in the expression of 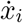 vanish since *x_ei_* = 0. Since *X_i_* is not a reactant in **R**_*j*_ this implies *β_ij_* > 0, *α_ij_* = 0. Therefore, *R_j_*(*x*) has a strictly positive coefficient, which implies 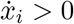 resulting in a contradiction.

Hence, we are ready to conclude the proof of Theorem 8:

##### Proof of Theorem 8-3).

By Lemma 3, there exists a steady state *x*^∗^ ∈ Ψ_*P*_ such that Γ*R*(*x*^∗^) = 0. Since dim(ker Γ) = 1, this implies that *R*(*x*^∗^) = *tv* for some *t* ≥ 0. Consider the case *t* = 0. This implies *R*(*x*^∗^) = 0. Then, 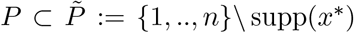. 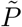 is a siphon by Lemma 4, and since 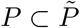 it is a critical deadlock. However, by Theorem 8-1), the network is does not admit a PWL RLF, which is a contradiction. If *t >* 0, this implies that *P* = ∅; giving a contradiction.

### 5 Concentration-dependent RLFs

#### 5.1 Proof of Theorem 9

Let 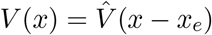. Then at those points *z* where 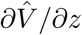 exists, we can write:

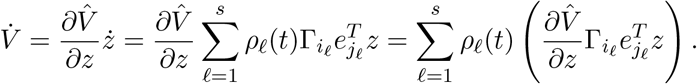

Since we have assumed that 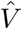 is a common Lyapunov function for the set of linear systems 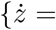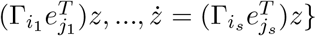, the proof can proceed in both directions in a similar way to the proof of Theorem 1. Notice that the constraint *D^T^z* = 0 is needed since 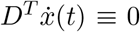 is implicit in the structure of the original system (1).

#### 5.2 Proof of Theorem 10

Positive definiteness is clearly satisfied. It remains to show the second condition. Let *z* = *x* − *x_e_*. Then, whenever 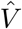 is differentiable:

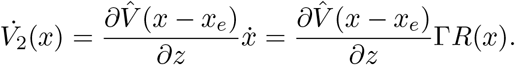

Before proceeding, we prove two statements: First, from (28), we get 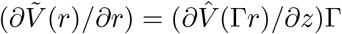. Second, note that *x* − *x_e_* ∈ Im(Γ), hence there exists *r* ∈ ℝ^*ν*^ such that Γ*r* = *x* − *x_e_*, where *r* can always be chosen nonnegative by assumption AS1. Hence, where 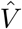 is differentiable, we can use (27) to write:

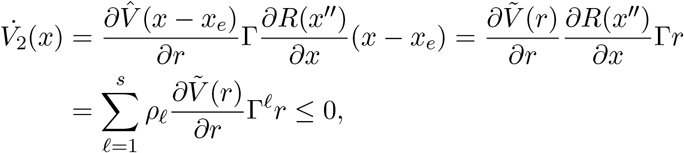

where the last inequality follows from (7). Lemma A1 implies that 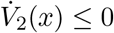 for all *x*.

#### 5.3 Proof of Theorem 11

The first statement follows from Theorem 10. In order to show the second statement, let 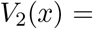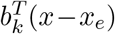, for 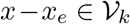. We will show that 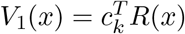, for each 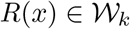 is nondecreasing along the trajectories. Without loss of generality, the partition matrix can be written in the form: 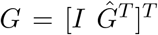. This representation implies that the sign of *x* − *x_e_* is determined in every region 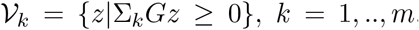, where Σ_*k*_ = diag[*σ_k_*_1_, …, *σkn*] are signature matrices. Now, assume that 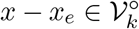. Then:

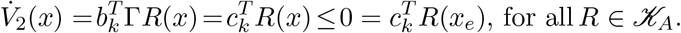

Let *R_j_*(*x*) ∈ supp *c_k_*, and let *α_ij_* > 0. Since *R* is nondecreasing by AK3, if 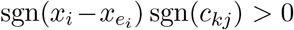, there exists 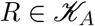 such that 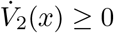. Hence, this implies that the inequality 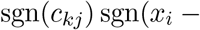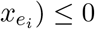 holds. Fix *j*. If there exists *i*_1_, *i*_2_ such that 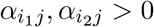 and 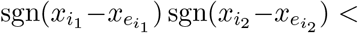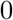, then *σ_kj_* ≔ 0. Otherwise, 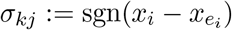 for some *i* such that *α_ij_* > 0.

Hence, in order to have 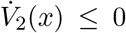 for all 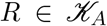 we need that 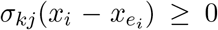 whenever 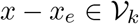, for all *k, j, i* with *α_ij_* > 0. By Farkas’ Lemma [13], this is equivalent to the existence of 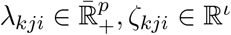, such that

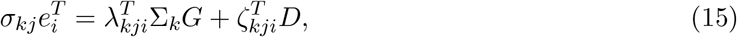

where *D^T^* ∈ ℝ^*ι*×*n*^ is a matrix whose columns are basis vectors for ker Γ^*T*^.

If we multiply both sides of (15) by Γ from the left, then we get condition C4 in [5, Theorem 4] which is necessary and sufficient for 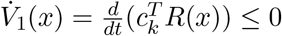.

### 6 Parameters for Figure 1

For the two mechanisms the total concentrations of the substrate and enzymes are [*X*_0_]_*T*_ = 6, [*E*]_*T*_ = 2.5, [*F*]_*T*_ = 6. The following ODE has been simulated for the distributive mechanism :

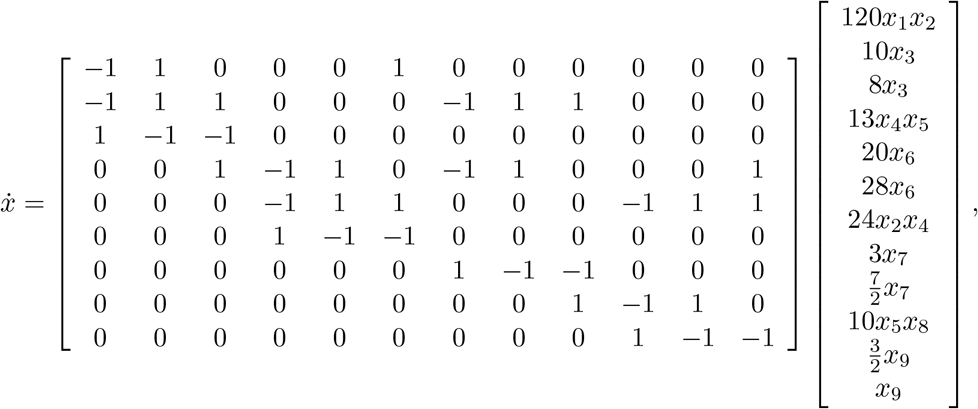

where *x*_1_ = [*X*_0_], *x*_2_ = [*E*], *x*_3_ = [*X*_0_*E*], *x*_4_ = [*X*_1_], *x*_5_ = [*F*], *x*_6_ = [*X*_1_*F*], *x*_7_ = [*X*_1_*E*], *x*_8_ = [*X*_2_], *x*_9_ = [*X*_2_*F*].

The following ODE has been simulated for the processive mechanism :

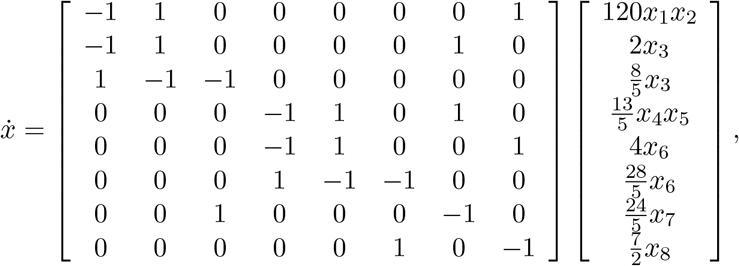

where *x*_1_ = [*X*], *x*_2_ = [*E*], *x*_3_ = [*X*_1_*E*], *x*_4_ = [*X*_2_], *x*_5_ = [*F*], *x*_6_ = [*X*_2_*F*], *x*_7_ = [*X*_2_*E*], *x*_8_ = [*x*], *x*_9_ = [*X*_1_*F*].

### 7 The Software Package LEARN

We describe the prerequisites of `LEARN`, the basic subroutines offered and few example runs. `LEARN` can be accessed at github.com/malirdwi/LEARN.

#### 7.1 Prerequisites

`LEARN` runs on MATLAB with the optimization and symbolic math toolboxes. Also, it needs the `cvx` package. The latest version of cvx is available on the link http://cvxr.com/cvx/download/. After download, the user must run `cvx_setup`. After `cvx_setup` reporting that cvx is successfully installed, `LEARN` should run without issues.

#### 7.2 List of Subroutines

The following subroutines are available. Note that all the subroutines below take Γ as an input which is the stoichiometry matrix of the network. If the network has an autocatalytic reaction then both matrices *A, B* need to be entered. (see the Methods section in the main text)

##### 7.2.1 Main subroutines

- `LEARNmain(Gamma)`: Prints a basic report on the network. This subroutine should be sufficient for most users. Examples will follow. Another parallel function, `LEARNmainplus(Gamma)`, is available which runs a more exhaustive RLF search.

##### 7.2.2 Basic subroutines

- `d=IsConservative(Gamma)`: Checks if the network is conservative. If it is, then the subroutine returns a positive vector 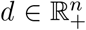 such that *d^T^* Γ = 0. If the network is not conservative then *d* returns a scalar 0.
- `v=IsAS1(Gamma)`: Checks if the stoichiometry matrix has a positive vector in its kernel. If it does, then the subroutine returns a positive vector 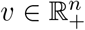 such that Γ*v* = 0. If the network is not conservative then *d* returns a scalar 0.
- `[flag,deadlock]=checkSiphons(Gamma)`: Checks if there are critical siphons and deadlocks. Each output can be either 0 or 1.
- `flag=checkMnetwork(Gamma)`: Checks if the network is an *M*-network. The output is either 0 or 1.

##### 7.2.3 Necessary Conditions

- `checkSiphonCondition(Gamma)`: Checks if the network violates the critical siphon necessary condition (Theorem 8). It prints a brief report.
- `flag=SignPatternCheck(Gamma)`: Checks if the network violates the sign pattern necessary condition [5, Theorem 9]. The output is either 0 or 1.
- `flag=checkPmatreix(Gamma)`: Checks if the network violates the *P* matrix necessary condition [5, Theorem 8]. The output is either 0 or 1.
- `flag=RobustNondegeneracy(Gamma)`: Checks if the network has a robustly non-degenerate Jacobian (Theorem 7). This only applies to networks that pass the *P* matrix test. The output is either 0 or 1.

##### 7.2.4 Construction of RLFs

- `C=ConstructGraphical(Gamma)`: Checks if the network admits the Max-Min RLF as given in Theorem 4. The output is *C*. If the method fails then *C* will be an empty matrix.
- `C=ConstructIterate(Gamma)`: Checks if the network admits an RLF as given in Theorem 3. The output is *C*. If the method fails then *C* will be an empty matrix.
- `[C,cvx]=ConstructLP(Gamma,H2,w,c)`: Checks if a non-autocatalytic network admits an RLF as given in Theorem 2. The last three inputs are optional. The output is *C* and the flag `cvx` to indicate that the RLF has been certified to be convex. The second input is *H*_2_ which are optional rows to add to the partitioning matrix *H* = Γ. The default value for *H*_2_ is an empty matrix. The third input is *w* and it is a flag to constrain the search to Sum-of-Currents RLFs. The default value is 1, but it is set to 0 in the `LEARNmainplus` subroutine. The fourth input is a flag to constrain the RLF to be convex. The default value is 0 which is the recommended value.
- `[C,cvx]=ConstructLPauto(A,B,H2,w,c)`: Checks if an *autocatalytic* network admits an RLF as given in Theorem 2. The remaining input structure is similar to the previous subroutine.
- `[C]=ConstructCoP(Gamma,H2)`: Checks if a non-autocatalytic network admits an RLF as given in Theorem 5. The last input is optional. The output is a tensor of PWQ RLF matrices. The second input is *H*_2_ which are optional rows to add to the partitioning matrix *H* = Γ. The default value for *H*_2_ is an empty matrix.

##### 7.2.5 Checking a candidate RLF

- `flag=CheckRLF(Gamma,C)`: Checks if 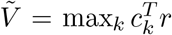 is an RLF for a non-autocatalytic network with the stoichiometry matrix Γ.

#### 7.3 Examples

All the examples are included in the folder `examples`.

##### 7.3.1 The double processive PTM cycle

This is the form of the input to `LEARN` for the network depicted in Fig. 9-b.

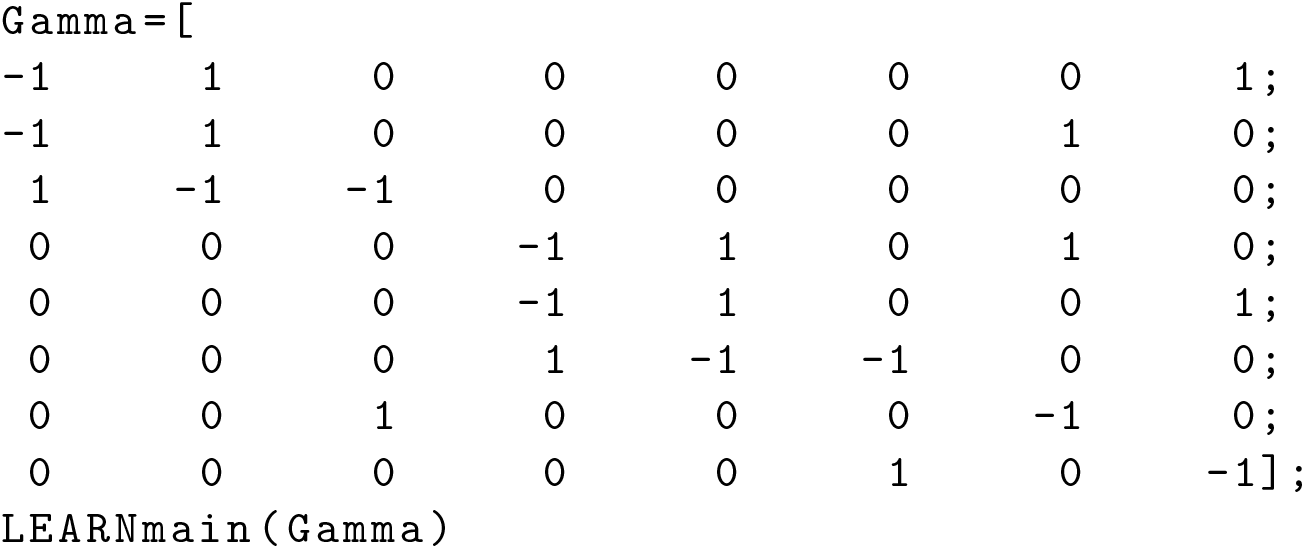

Note that the stoichiometry matrix Γ can be easily written from a list of reactions. The output of `LEARN` is as follows:

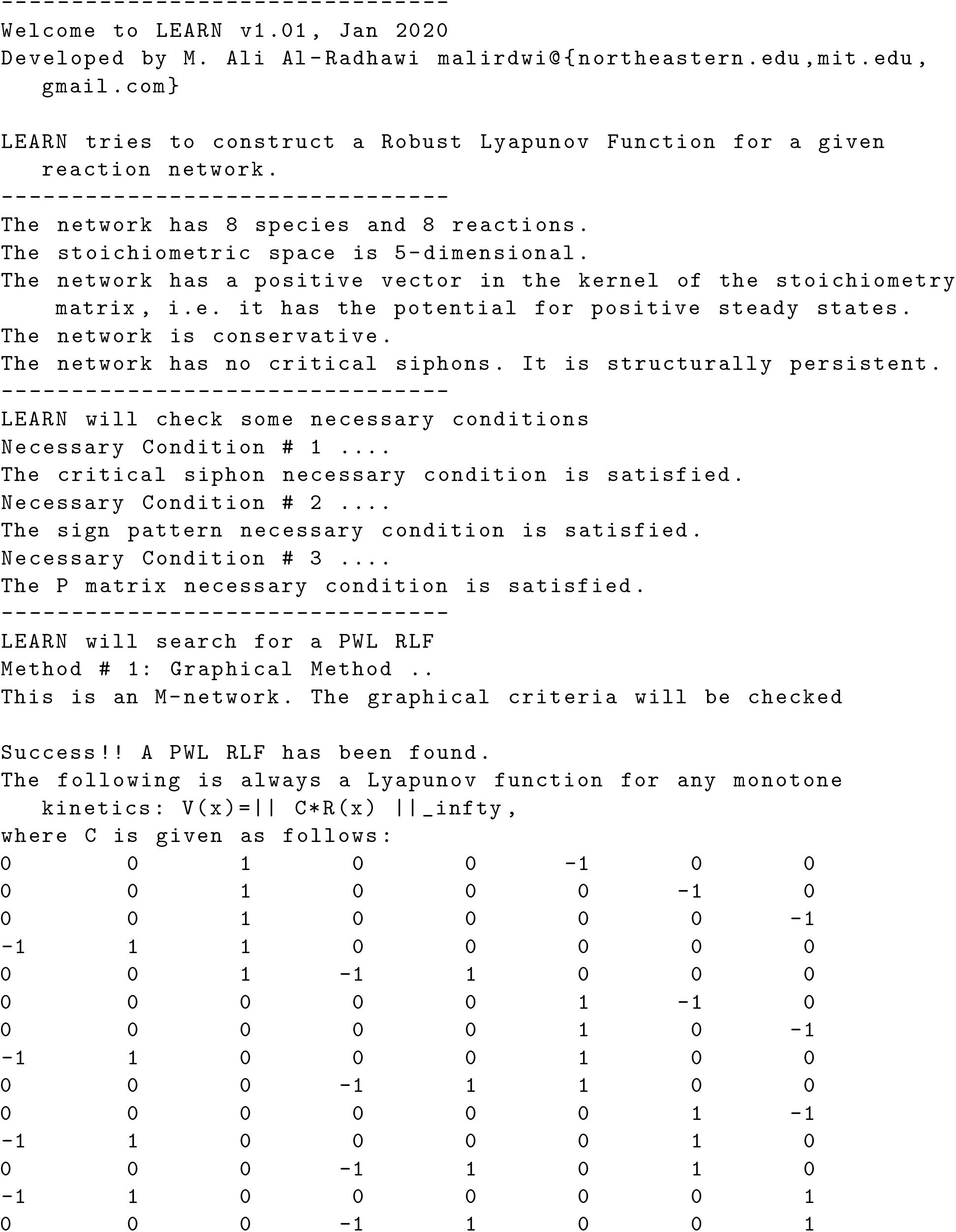

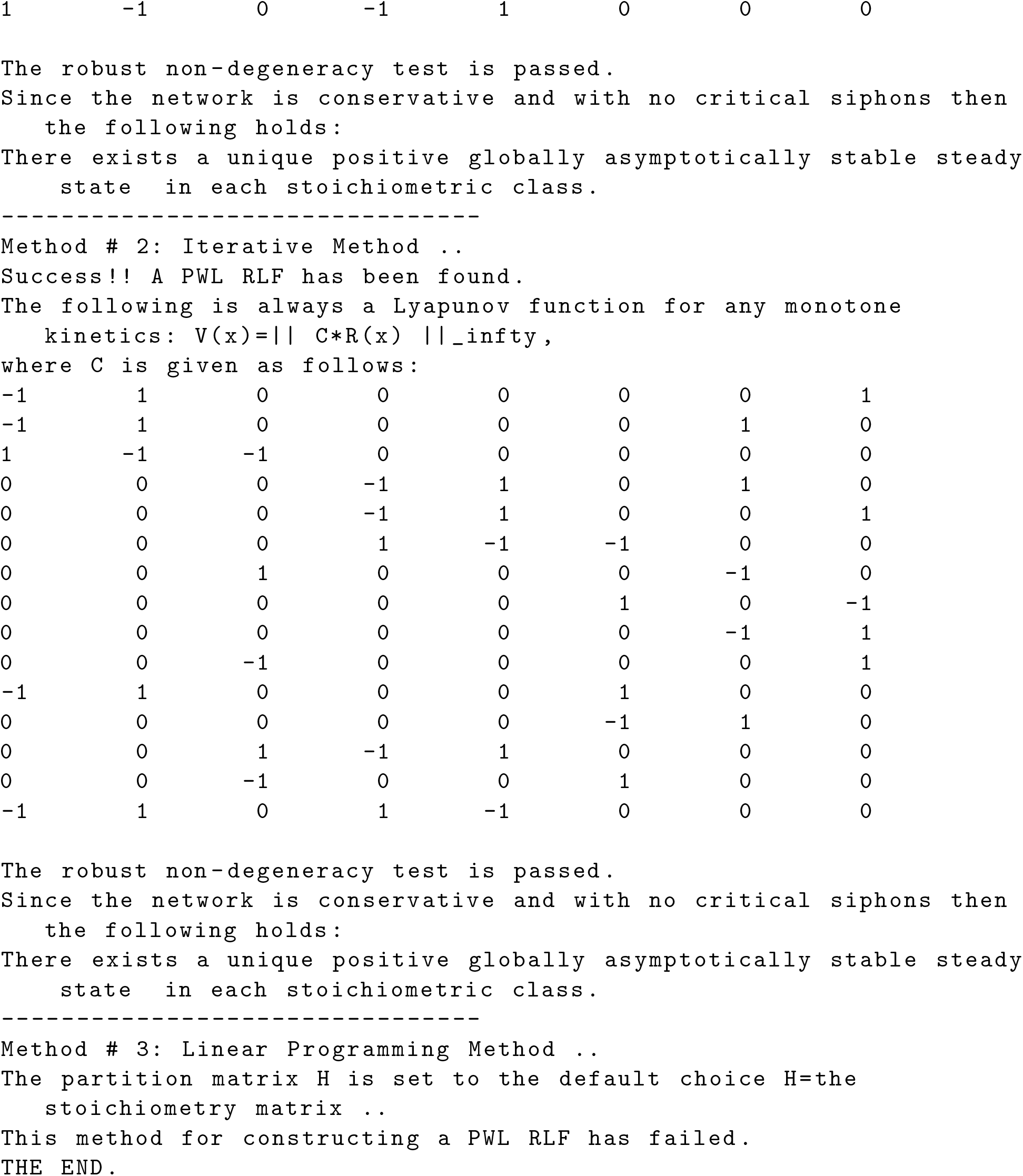

##### 7.3.2 The double distributive PTM cycle

This is the output of `LEARNmain` for the network depicted in Fig. 9-d.

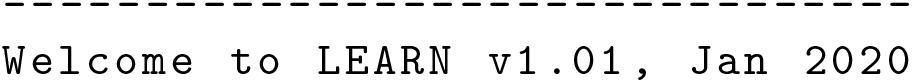

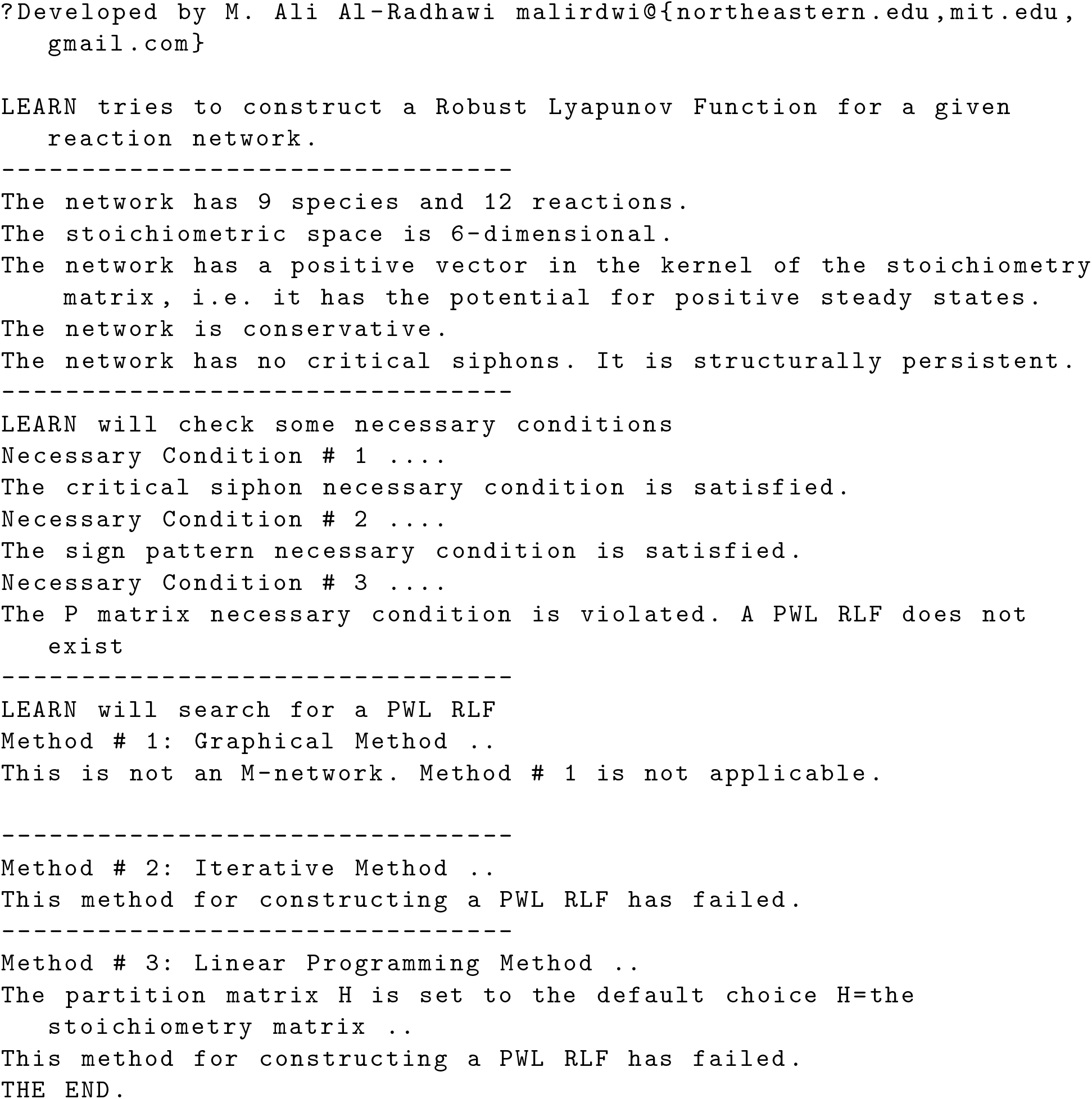

##### 7.3.3 The McKeithan Network

This is the output of `LEARNmain` for the network depicted in Fig. 11-a with *N* = 2.

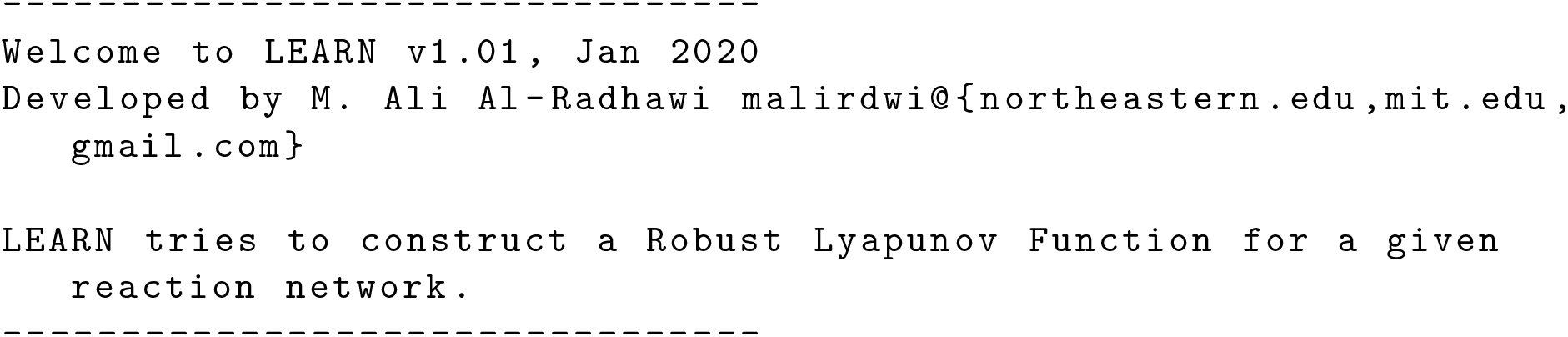

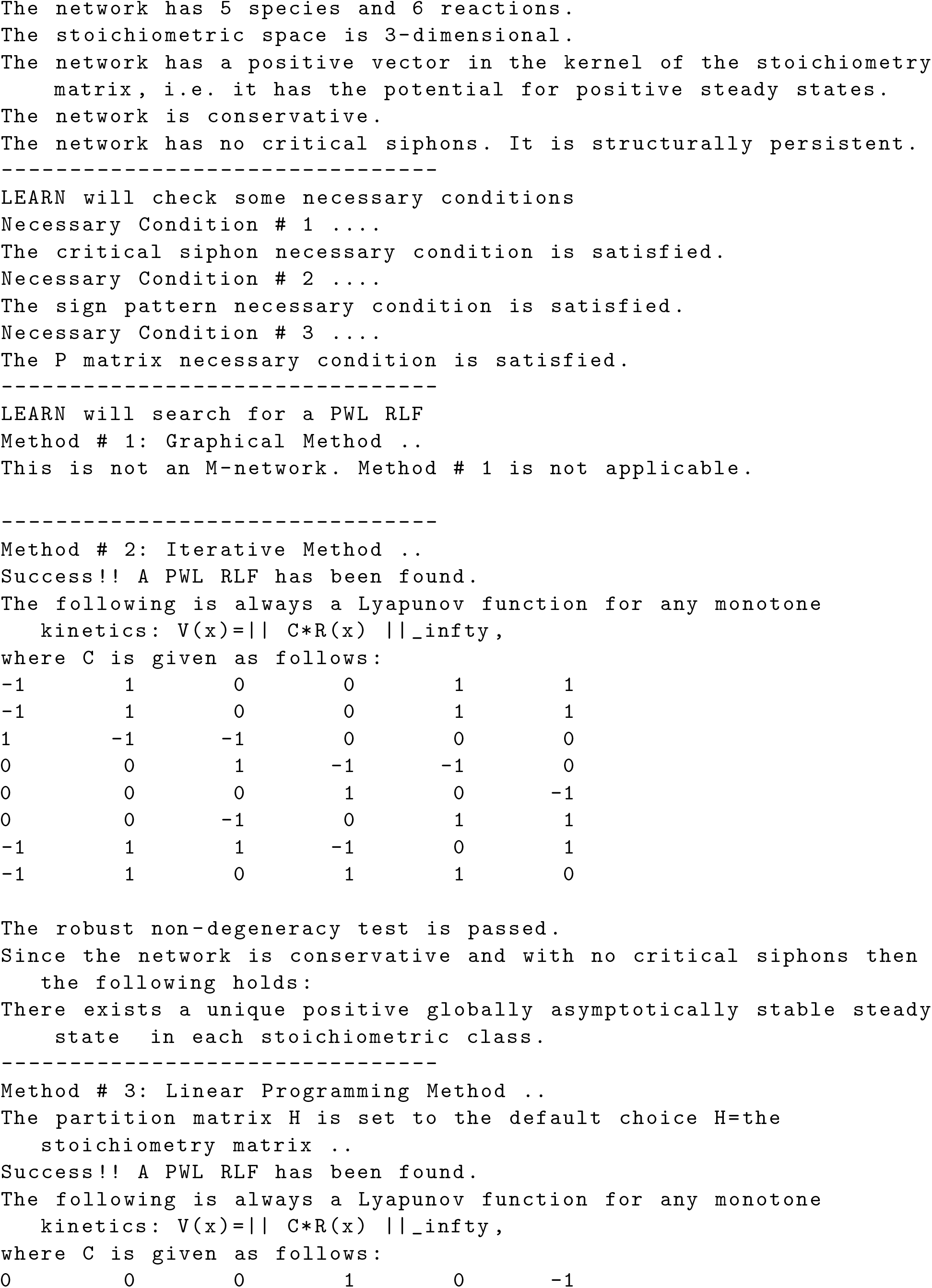

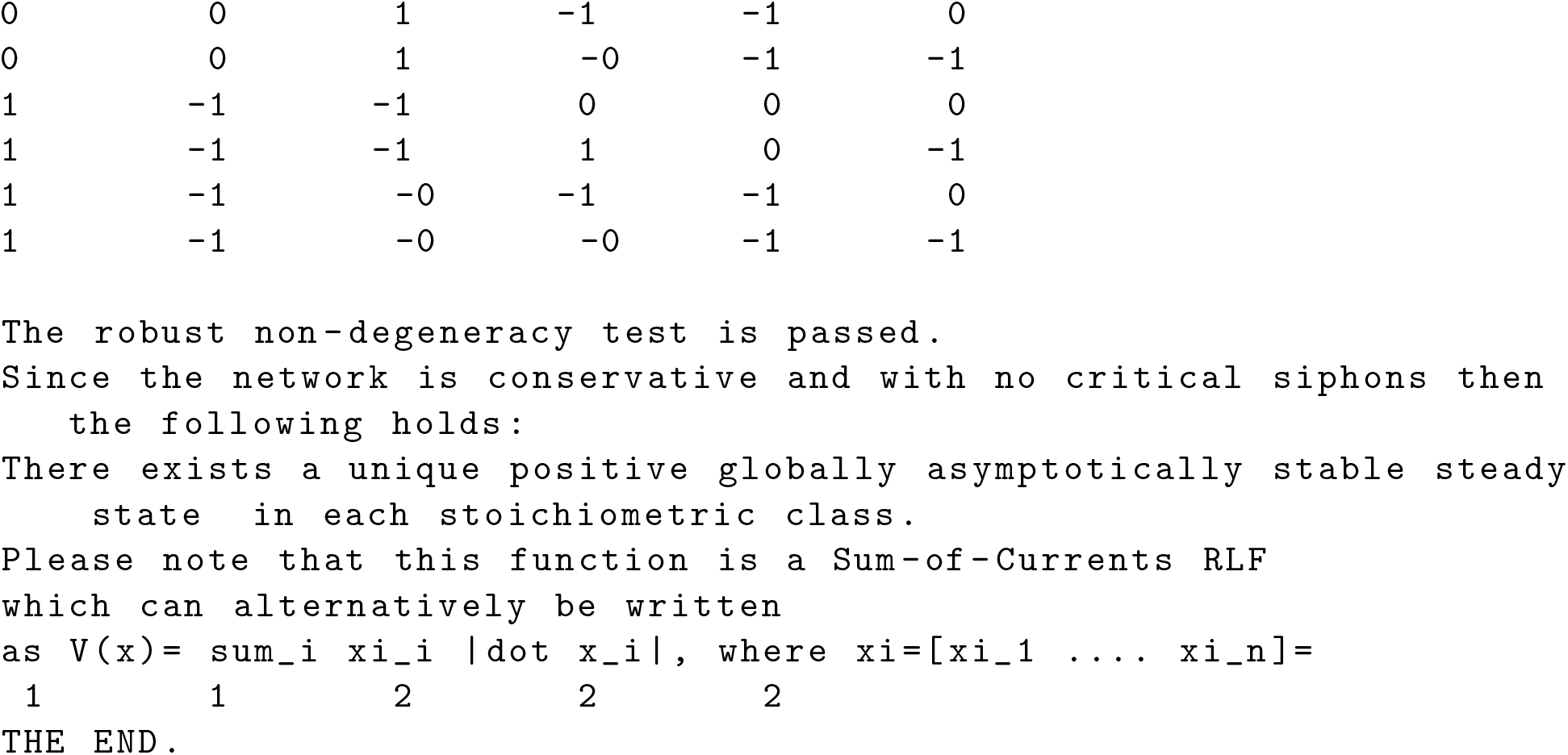

## Notes

http://www.github.com/malirdwi/LEARN

